# Cell competition overcomes host tissue resistance to unleash tumour growth in a *Drosophila* brain cancer model

**DOI:** 10.1101/2025.08.05.668787

**Authors:** Marco Gualtieri, Saline Jabre, Damien Mornico, Maëlwenn Hamon, Cédric Maurange, Pauline Spéder

**Affiliations:** Institut Pasteur, CNRS, UMR3738, Université Paris Cité, Equipe labellisée Fondation ARC pour la Recherche contre le Cancer Paris, France; Sorbonne Université, Collège Doctoral, ED515, Paris, France; Hub de Bioinformatique et Biostatistique - Département Biologie Computationnelle, Institut Pasteur, USR 3756 CNRS, Paris, France; Aix Marseille University, CNRS, IBDM, Equipe labellisée Ligue Contre le Cancer, Marseille 13009, France

## Abstract

Primary tumours of the central nervous system are extremely aggressive and often incurable. While tumours are known to interact with their microenvironment, how the interplay between tumour cells and the host tissue controls tumour progression remains elusive. We addressed this question in a *Drosophila* model of cancer stem cell-driven tumour which originates during development and grows extensively within a network of cortex glia cells (CG) through adulthood. We found that tumour growth induces progressive remodeling of the CG network and leads to CG apoptosis through cancer stem cell-driven competition, a process which in turn unleashes tumour growth. Notably, preventing CG death reduces tumour growth, revealing a resistance capacity within the CG. Transcriptional profiling of CG cells at different stages revealed a biphasic response, featuring an early specific signature followed by a later collapse of essential cellular functions and pathways, including the conserved JNK pathway. JNK signalling in CG normally plays a neuroprotective role and initially hinders tumour progression, providing resistance. However, its subsequent downregulation eventually brings the collapse of the host tissue, ultimately curbing tumour growth. This study uncovers a dynamic and complex interplay between host tissue resistance and tumour-driven competition, which shapes tumour progression.

## Introduction

Primary tumours of the central nervous system (CNS) are extremely aggressive tumours which originate from and develop within the brain or spinal cord (*1*, *2*). They are still extraordinarily difficult to treat despite combined surgery, chemotherapy and radiotherapy, owing to their infiltrating nature challenging resection and to their place of origin, shielded behind strong cellular barriers and made of poorly replaceable cells. Survivors are left with important neurological sequelae and a degraded quality of life, from the tumour and/or the treatments. Tumour recurrence is also high. Another specificity of CNS tumours is their high incidence in children, accounting for 26% of new cancer cases versus 1.5% in adults (*3*) and making them the leading cause of cancer mortality during childhood. In contrast to adult tumours, pediatric tumours do not seem to accumulate multiple oncogenic mutations, while sharing highly proliferating and invasive properties (*4*).

CNS tumours originate from and are maintained by a restricted number of highly proliferative cells at the apex of the tumour cellular hierarchy. These cells, identified in several primary CNS tumours (*5–9*), are coined cancer stem cells (CSCs) and share many features of neural stem cells (NSCs), the multipotent progenitors of the CNS, whose transformation can lead to cancer development (reviewed in (*10–13*)). CSCs display NSC molecular markers, exhibit self-renewal potential and can birth differentiated progeny, while supporting the malignant tumour population through uncontrolled amplification. Developmental signalling pathways present in NSCs also re-emerge in CSCs to promote growth and invasiveness (*13*, *14*). CSCs are at the apex of the tumour cellular hierarchy, can re-initiate the tumour (*15*) and are proposed to resist current treatments, supporting cancer recurrence (*16*). CSC properties are increasingly linked to tumour growth and multi-scale complexity (competition, composition, invasiveness) (reviewed in (*10*, *14*, *17*)).

While tumour growth is fueled by the intrinsic proliferative ability of CSCs, it is now known that it can be dramatically influenced by extrinsic factors. Tumours indeed develop within, both with and against, the host tissue. The tumour microenvironment (TME) builds from the host components in vicinity to the tumour, and evolves over time, resulting from the progressive remodelling of the host tissue as well as from the contribution of CSC-produced cells. The interplay between tumour cells and the host tissue/TME is complex and multifaceted, and appears as a crucial driver of cancer progression (*18*, *19*). In CNS tumours, the host tissue, and thus the TME, is particularly intricate and made of highly diverse cellular components of various origins, including neurons, glial cells, the blood-brain barrier and resident immune cells (*20–22*).

Hijacking and rewiring of host cell functions by tumour cells have been well documented across various cancer types, including CNS tumours, to support tumour progression by conferring metabolic and immune advantages (*23*, *24*). For instance, cross-talk between CSCs and endothelial cells increases angiogenesis and accelerates CSCs’ self-renewing capacity (*25*). Tumour cells can actually set up connected networks between themselves or with host cells, a property linked to tumour amplification and resistance to treatment (*26*). Moreover, neuron-tumour interactions influence tumourigenesis at multiple steps and through diverse mechanisms (reviewed in (*27*, *28*)), including direct interaction through synapses (*29*) and coopting neuronal migration (*26*). Glial cells, namely oligodendrocytes and astrocytes, are also part to the TME, yet their interactions with tumour cells are still poorly characterized (reviewed in (*30*, *31*)). Tumour reactive-astrocytes promote glioma invasion through different factors (*31–34*). Interestingly, in early stages, astrocytes seem to exert anti-tumour, cytotoxic effects before becoming reactive, highlighting a dynamic interplay between the glial TME and tumour cells (*35*). Despite growing attention to the role of the host tissue in oncogenesis, and evidence that their properties are remodelled in cancer (*36*, *37*), the contribution of glial cells to tumour progression is still little understood.

Tumours can also reshape the TME by removing cells and have been proposed to use cell competition to create physical and functional space for invasion (*38*). Cell competition has been defined as a context-dependent process where one cell population (“winner cells”) induces the apoptosis of another (“looser cells”), leading to its elimination (*39*). This mechanism is thought to support the proliferation of rapid-growing cells (known as super-competitors) by removing and replacing neighbouring healthy cells. As such, this concept, originally investigated in development and discovered in *Drosophila* epithelia (*40*, *41*), has important implications for cancer. A number of studies in epithelial tumours have now shown that tumour cells can kill their neighbouring cells, either less fit tumour clones (thus shaping tumour clonality) or healthy host cells (*42–46*), to boost their growth (reviewed in (*47–49*). Whether similar mechanisms operate in the non-epithelial CNS tumours remain poorly characterized. Clonal competition happens in glioblastoma, influencing tumour genetic composition (*50*). Moreover, CNS tumours grow in a dense tissue with strong boundaries, creating solid stress which induces neuronal death (*51*). High levels of cell death are detected across tumours and linked to poor prognosis, with increasing evidence showing that dying cells positively affect tumour cells by releasing beneficial factors (*52*, *53*). However, while cell death regulation in tumour cells has been extensively studied, given its potential for therapeutic targeting and its role in tumour resistance (*54*, *55*), the extent and function of cell death in the different host cells of the CNS, and whether their sensitivities change in tumour conditions, remain poorly known (*56*). In particular, whether *bona fidae* cell competition between cancer cells and the host tissue happens in CNS tumours, and its relevance for the pathology, is not known.

Here, we used a *Drosophila* model of CNS tumour of developmental origin to decipher the dynamics between the host tissue, specifically glial cells, and tumour progression. Aggressive, CSC-driven tumours can be induced from transformed NSCs during the developing larval stage, resulting in fast-growing tumours in adults. During normal development, Type I NSCs, the most prominent NSC subtype in the fly CNS outside of the visual system, divide asymmetrically to self-renew while generating an intermediate progenitor cell called ganglion mother cell (GMC), which divides once more to produce two neurons or glial cells (**Fig. 1A**, left panel) (*57*). Individual NSCs can be dysregulated in a cell-autonomous fashion to generate tumours (*58*, *59*). In particular, the loss of the Prox-related homeobox transcription factor Prospero (Pros) produces GMCs which fail to differentiate and revert to a stem cell-like state (*60*) (**Fig. 1A**, right panel). Each *pros^-^*NSC gives rise to a clonal tumour lineage mainly composed of NSC-like cells, which amplify at the expense of normal neuronal output and which persist and develop in the adult stage of the fly (**Fig. 1B-C**). To form tumours, *pros^-^* NSCs must be induced during an early developmental window when the temporal factors Chinmo and Imp are expressed (**Fig. 1A**, insert) (*61*). These *chinmo^+^ Imp^+^ pros^-^* NSCs have high proliferative potential and persist indefinitely in the tumour, acting as self-renewing CSCs (*62*). They can also generate more restricted *Syp^+^ pros^-^* progenitors in which *chinmo* and *Imp* are silenced following the emergence of the RNA-binding protein Syp, leading to limited proliferative capacity. Together, these populations form a heterogeneous, hierarchically organized tumour with *Chinmo^+^ Imp^+^* CSCs at the apex (*62*) (**Fig. 1A**). Chinmo and Imp are reminiscent of oncofetal genes, whose expression is normally restricted to embryonic/fetal stages and reactivated in tumours. They appear to promote tumour growth by boosting cell metabolism and self-renewal. These hierarchical CSC-driven tumours, induced during developmental stages by the inactivation of a single gene, through co-option of developmental programs, offer a simplified yet relevant model for studying pediatric tumour progression (*59*, *61–63*).

**Figure 1.**
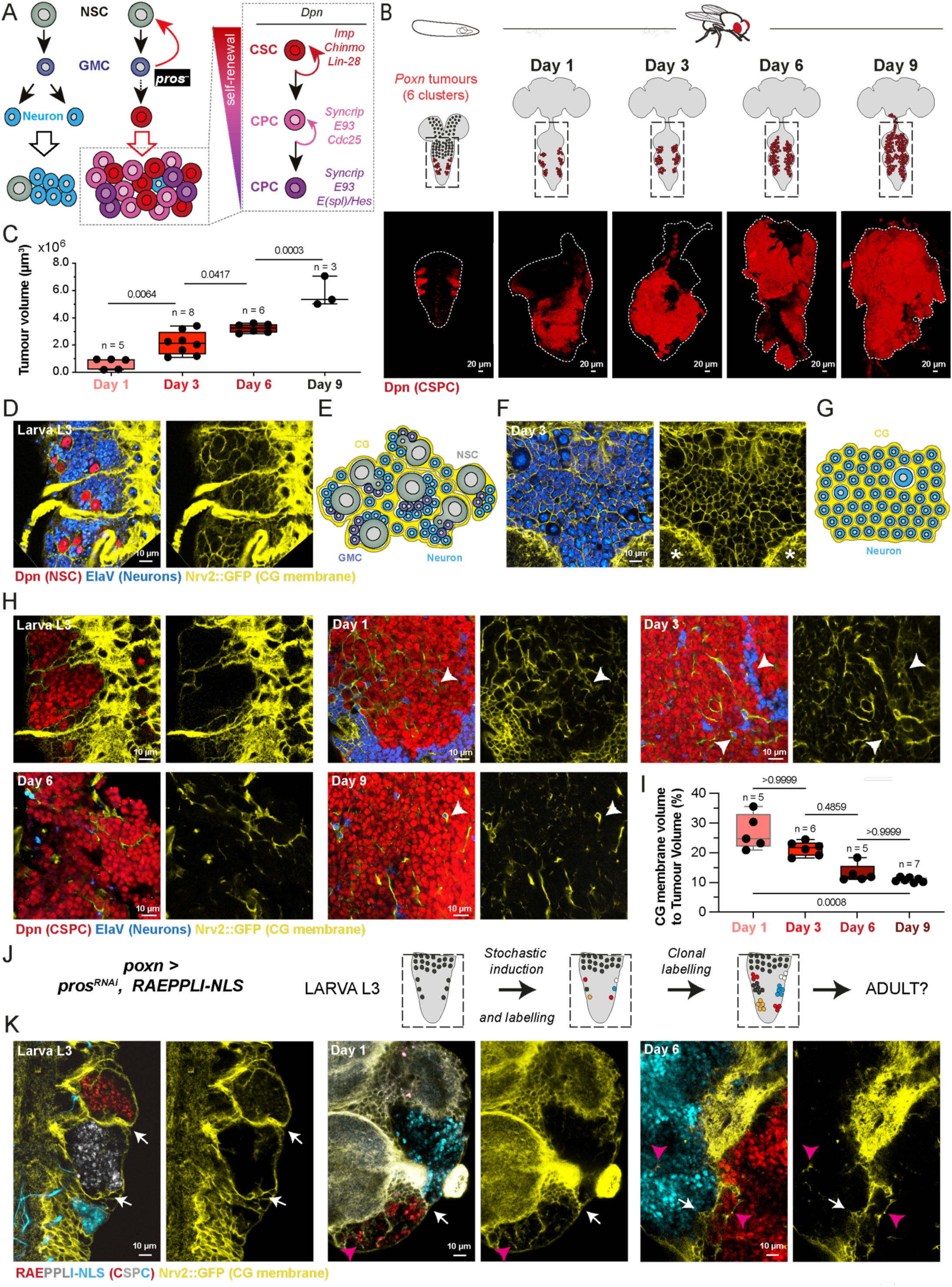
Cancer stem cell-driven tumours remodel the host glial microenvironment (A) Schematic of tumour initiation from transformed neural stem cells. During normal larval neurogenesis, neural stem cells (NSCs, grey) divide asymmetrically to generate one ganglion mother cell (GMC, purple), a precursor which will divide further only once in a symmetric fashion to produce two neurons (N, blue, depicted here) or more rarely glial cells. Upon the loss of *prospero* (*pros*), a gene driving NSC differentiation, GMCs revert to a stem cell-like fate, transforming into Cancer Stem Cells (CSCs, red). These CSCs (marked by the expression of Imp, Chinmo and Lin-28) are at the apex of the cellular hierarchy (insert). They will generate the diversity of the tumour, composed of CSCs and Cancer Progenitors Cells (CPCs, in pink and purple shades) of varied potential (marked by Syncrip and E93, and differentiated by the expression of Cdc25 or E(spl)/Hes), and occasionally neurons. All CSCs and CPCs (collectively refered as to CSPCs) express the transcription factor Deadpan (Dpn). (B) Schematic and illustration of tumour growth in *pros^poxn^* tumours (*poxn > pros^RNAi^).* Six individual tumours are initiated at the larval stage by the specific knockdown of *pros* in the six Poxn^+^ NSCs located in the ventral nerve cord (VNC, highlighted in dashed black rectangles), in a dorsal and posterior thoracic fashion, using a restricted GAL4 driver (*poxn-GAL4*). Shortly after adult eclosion (day 0), the tumours have started growing (day 1) and they will keep expanding towards the central brain until around day 9 where the flies die. Confocal pictures are 3D reconstructions of tumours stained with Deadpan (Dpn, red), a marker of NSCs and CSCs. Dashed white lines delineate the VNC. (C) Quantification of tumour volume over time. n = VNC. One-way ANOVA with Tukey’s multiple comparison tests. (D) Confocal image showing the NSC niche at third larval stage (L3), in which Poxn^+^ NSCs transform into CSCs upon *pros* knockdown. NSCs and their neuronal progeny are closely enwrapped by cortex glia (CG) membranes. (E) Schematic of the NSC niche at larval stage. (F) Confocal image showing the host cellular microenvironment, in which the CSC-driven tumour will expand, at adult stage. Neurons are surrounded by a tight meshwork of CG membranes. Neuropile structures, in which synapses form, are indicated by white stars. (G) Schematic of the host cellular microenvironment at adult stage (H) Crops of confocal images illustrating the progressive remodelling of CG membranes in the VNC upon tumour progression. White arrowheads indicate examples of individually enwrapped neurons. (I) Quantification of CG membranes volume compared to tumour volume along tumour progression. n = tumour. Kruskal–Wallis H test with Dunn’s multiple comparisons test. (J) Principle of clonal tumour labeling using the genetic construct Raeppli-NLS. Early induction by heatshock at larval stage leads to the labelling of each of the six original Poxn^+^ CSCs by one out of four possible colour choices. The entire lineage of each CSC will inherit the colour which will be genetically maintained over the entire lifespan, thus allowing the tracking of the six individual tumours within the CNS. (K) Confocal images of *pros^poxn^* tumours labelled by Raeppli-NLS within the CG membrane (yellow) at third larval stage (L3) and days 1 and 6 of adulthood. While the CG remodels, the entire tumour lineages keep together and stay separated from one another by a layer of CG membranes, similarly to larval NSC lineages. White arrows show the location of the CG separating tumour clones. Pink arrowheads indicate examples of CG membranes infiltrating within the tumour mass. For box plots, individual values are superimposed. See also Table S1.

These *pros^-^* CSC-driven tumours develop within a host tissue made of diverse cell types making up the NSC niche (*64*), including the NSCs themselves, neurons, multiple glial subtypes (*65*) and a blood-brain barrier (*66*) (**Fig. 1D-E**). The blood-brain barrier and glia provide lifelong trophic support and regulatory inputs to NSCs and neurons. In particular, the cortex glia (CG) form a highly connected membrane meshwork (*67–71*), infiltrating in-between individual NSC lineages and neurons. The CG support and protect NSCs (*72–75*) and are essential for neuronal positioning and survival (*67*, *69*, *76–78*). In the adult, the CG network exists around neurons (**Fig. 1F-G**) and support their functions, such as sleep regulation (*79*, *80*). Notably, the CG structure, composed of interconnected cells which communicate and maintain close contact with neurons, resembles the astrocytic networks found throughout the brain (*81*). Moreover, CG interface with the blood-brain barrier, akin to the mammalian *glia limitans*, another reticular structure made by astrocytic endfeet (*82*).

Here, we found that CG cells initially remodel their morphology to infiltrate the tumour, and provide resistance to tumour growth. However CSC-mediated cell competition takes place and results in the elimination of CG cells by apoptosis, a phenomenon which boosts tumour growth. This biphasic TME remodelling is mirrored by transcriptional changes in CG cells, characterized by an early specific signature followed by the collapse of core cell functions. Amongst these changes, the downregulation in the CG of the normally active and neuroprotective JNK pathway first helps tumour growth but later hinders it, a shift coinciding with neuronal loss and overall tissue breakdown, and implying that JNK morphes from an anti- to a pro-tumour role during the course of the tumour. These findings highlight the complexity of tumour and host tissue interactions, and identify both cell competition and glial cells as crucial players in CNS tumours.

## RESULTS

### Cortex glia architecture is remodelled during tumour progression

We used the UAS/GAL4 system to knockdown *pros* by driving a specific RNAi under the genetic control of a Paired-box transcription factor Pox-Neuro (*poxn*) enhancer that is active in only six Type I NSCs in the larval ventral nerve cord (VNC) (*83*) This system allows the generation of six trackable tumours (hereafter referred to as *pros^poxn^* tumours) in stereotyped locations in the larval VNC (**Fig. 1B**) (*61*). Their cancer stem and progenitor cell population (herein called CSPC), which makes most of the tumour bulk, can be marked with NSC determinants such as the transcription factor Deadpan (Dpn) (**Fig. 1A**, insert). These *pros^poxn^*tumours grow from larval stages. A large part of CSPCs differentiate during metamorphosis, in response to hormonal pulses, but a fraction persists in adult. From adult eclosion (day 0), the persisting fraction of CSPCs continues to overproliferate, covering the entire VNC after a few days, further invading the central brain, and ultimately killing the fly around days 10-12 (**Fig. 1B-C** and (*61*)).

To understand the interplay between CG cells and tumour progression, we first assessed their morphology. During normal development, NSC lineages are found embedded in a membranous chamber made by the CG (**Fig. 1D-E**), a grid-type pattern maintained by a balance of timely set-up differential adhesions (*78*). This rule of whole lineage encasing stops when neurons mature, and become individually enwrapped (*68*, *78*). Previous work also showed that NSC-derived tumours of different origins, induced in early developmental stages, stay confined within a continuous CG enclosure during their growth at larval stage (*78*). Using the *Nrv2::GFP* knock-in line to visualize the CG membranes and Dpn staining to detect the CSPCs, we found that each of the six *pros^poxn^* tumours similarly stayed contained inside CG chambers during larval stages (**Fig. 1H and S1A**, L3 larval stage), with the tumour clones not merging with each other. The individual tumours were also devoided of internal CG membrane.

We wondered whether such embedding was maintained during tumour progression in adulthood and how effectively CG coped with the dramatic tumour growth. We found that, in most VNCs, individual *pros^poxn^* tumours were not no longer distinguishable by day 1, and we rather observed one tumour mass spread throughout the whole VNC and infiltrated by discontinuous CG membranes (**Fig. 1H and S1A**, Days 1, 3, 6 and 9). This was accompanied by a progressive remodeling of the CG structure along tumour progression, displaying altered membrane morphology and eventually losing its seemingly continuous network organization. Quantifying the ratio between CG membrane and tumour volumes revealed an overall decrease along tumour progression (**Fig. 1G**). Notably, neurons continued to be individually encased by CG membranes, even the rare ones within the tumour mass (**Fig. 1H**, white arrowheads).

While the CG structure was disorganised, making individual tumours undiscernible, it remained to be addressed whether the six *pros^poxn^* tumours were meeting each other and truly merging or whether they stayed separated from each other by CG membranes. To distinguish the original six tumour clusters in the adult, we took advantage of a multicolour clonal labelling technique, Raeppli (*84*), in which the stochastic expression of one out of four fluorophores, targeted to the nucleus by a nuclear localization sequence (NLS-Raeppli), can be restricted to Poxn^+^ NSC lineages through the use of the GAL4/UAS system. Induction is done at larval stage to label the Poxn^+^ NSCs from which the tumours originate from (**Fig. 1J**). Ultimately, in the tumour condition, each of the six initial NSCs generates a tumour clone stably marked by one out of the four possible fluorophores (**Fig. 1K** and **S1B**, Larval stage) and which can be tracked throughout adult stage (**Fig. S1C**). First, we found that tumour clones do not seem to mingle, but rather stay on their own, as a compact group of same colour cells (**Fig. 1K** and **S1B**, days 1 and 6), a behaviour similar to what was observed for these and other tumours in larval stages. Focusing on the zone between two clones, we observed that CG membranes, labelled by Nrv2::GFP, were still present between different *pros^poxn^* tumours at later stages (days 3 and 6, white arrows), suggesting that the CG still maintain its singular embedding of clonal lineages. We however noticed the presence of many infiltrating CG membranes within individual *pros^poxn^*tumours (**Fig. 1K** and **Fig. S1D**, pink arrowheads), confirming our previous observation.

Altogether, these data suggest that during tumour progression, CG maintain the developmental principle of encasing whole lineages for undifferentiated cells and encasing individual cells for neurons. Yet, CG organisation remodels around tumours, with the early formation of internal membrane projections infiltrating in-between CSPCs and whose density within the tumour decreases overtime.

### Cancer stem cell-driven cell competition induces close-by cortex glia apoptosis

The dramatic expansion in volume of *pros^poxn^* tumours, together with CG remodelling, made us wonder how they could push against the host tissue and, specifically, whether it was at the expense of neurons and glia.

We first asked whether host cells were displaced or lost during tumour progression. We stained tumour and control CNSs at different stages of tumour progression (3 and 6 days old adults) with a marker for glial (transcription factor Repo) and neuronal (transcription factor ElaV) fates. We first found that both glia and neuron nuclei appear mislocalized in the tumour VNC compared to control (**Fig. 2A**), exhibiting patterns ranging from strewn to clumped throughout tumour progression. We also noticed a qualitative decrease in the overall glial staining. We then quantified the neuronal and glial populations using the ElaV and Repo markers. Since individual neuronal signal was challenging to segment (due to high neuronal density), we decided to use the total volume of nuclei over time as a proxy volume of the cell population for each cell type (see Methods). Thus, from now on, neuronal or glial volume will refer to total nuclei volume for the respective population. Quantification of neuronal volume showed no significant difference between control and tumour condition (**Fig. 2B**). In contrast, glial volume was significantly reduced in tumour conditions compared to controls, at both day 3 and day 6 (**Fig. 2C**). These results show that tumour progression leads to glial, but not neuronal loss, suggesting that neurons are more resistant to tumour progression or/and that the tumour primarily affects glial cells.

**Figure 2.**
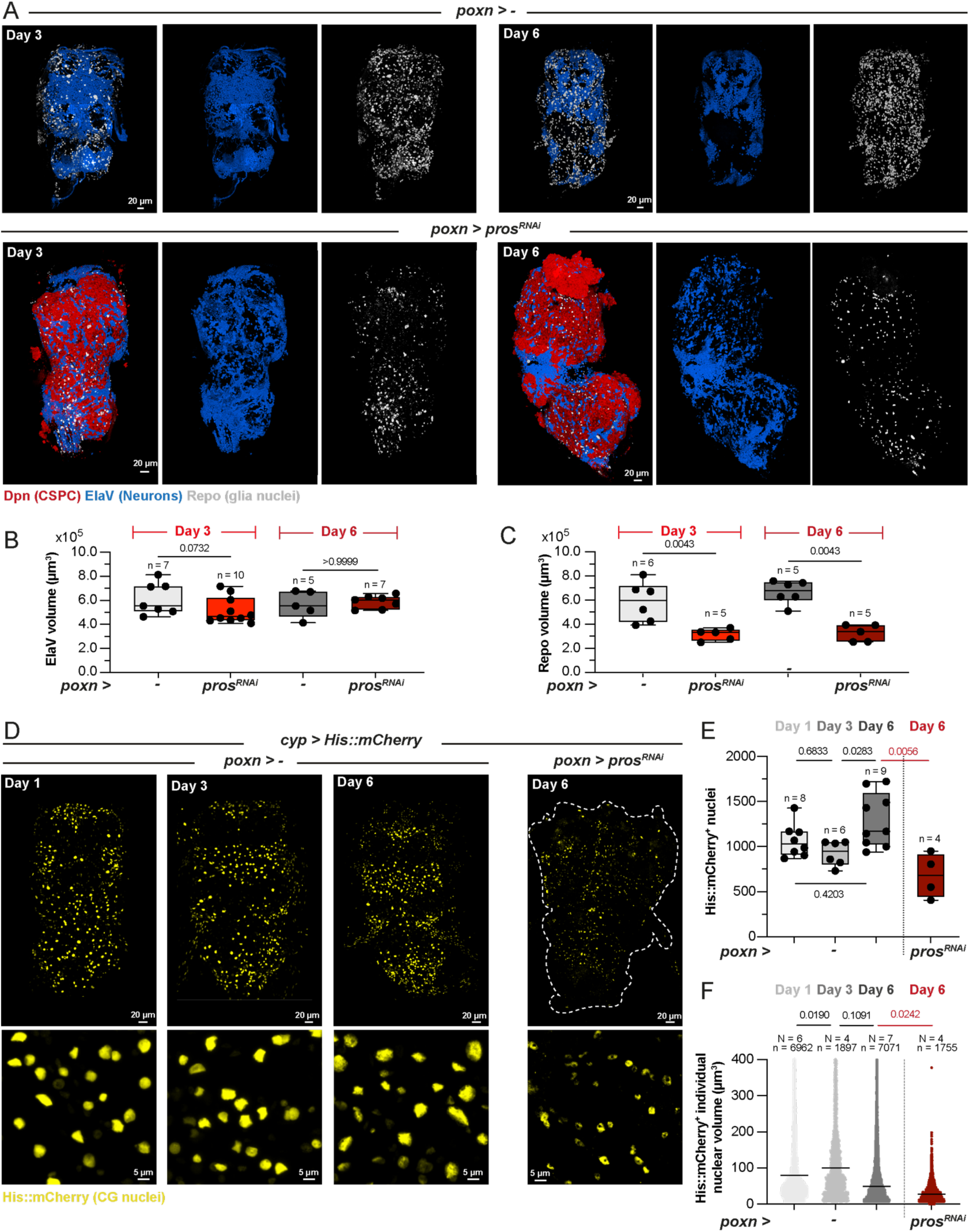
Cortex glia, but not neurons, are lost upon tumour progression. (A) 3D reconstruction of confocal images showing whole VNCs stained for neuronal (ElaV, blue) and glial (Repo, gray) nuclei for control (*poxn > -*) and *pros^poxn^* tumour conditions (*poxn > pros^RNAi^*) at days 3 and 6 of adulthood. (B) Box plot of total neuron (ElaV^+^) volume for control (*poxn > -*) and *pros^poxn^* tumour conditions (*poxn > pros^RNAi^*) at days 3 and 6 of adulthood. n = VNC. Mann-Whitney U tests for each day. (C) Box plot of total glia (Repo^+^) volume for control (*poxn > -*) and *pros^poxn^* tumour conditions (*poxn > pros^RNAi^*) at days 3 and 6 of adulthood. n = VNC. Mann-Whitney U tests for each day. (D) Confocal images of CG nuclei genetically labelled by a His::mCherry fusion (*cyp > His::mCherry*, yellow) for control (*poxn > -*) for days 1, 3 and 6 of adulthood and *pros^poxn^*tumour conditions (*poxn > pros^RNAi^*) for day 6 of adulthood. Upper panels show 3D reconstruction of whole VNCs and lower panels show a close-up projection. (E) Box plot of CG nuclei number for control (*poxn > -*) at days 1, 3 and 6 of adulthood and *pros^poxn^* tumour conditions (*poxn > pros^RNAi^*) at day 6 of adulthood. n = VNC. One-way ANOVA with Tukey’s multiple comparisons test between the control conditions. Mann-Whitney U tests between chosen pairs. (F) Dot plot of CG individual nuclear volume for control (*poxn > -*) at days 1, 3 and 6 of adulthood and *pros^poxn^* tumour conditions (*poxn > pros^RNAi^*) at day 6 of adulthood. Dot = CG nucleus. N = VNC. n = CG nucleus. Mann-Whitney U tests between chosen pairs were performed on the mean values per VNC. For box plots, individual values are superimposed. See also Table S1.

Glial loss in tumour conditions could result either from disruption of physiological, programmed proliferation or from increased cell death. While the former explanation appeared less likely, as glial cells are deemed post-mitotic, it remained to be ruled out. Moreover, whether the glial loss could be allocated, at least in part, to the CG, was also left to be shown. To address both points, we specifically labelled CG nuclei using a *His::mCherry* transgene under the control of a CG-specific promoter (*cyp4g15*) (**Fig. 2D**), and quantified both the number and individual volume of CG nuclei at days 1, 3 and 6 in normal and tumour condition (**Fig. 2E-F**). Unexpectedly, we first found that in normal situation, CG displayed stereotyped and timed changes in nuclei number and individual nuclear volume over time, suggestive of some proliferative behaviour. CG nuclei number stayed at similar level between day 1 and day 3 but showed a distinct increase at day 6 (**Fig. 2E**), while CG individual nuclear volume increased between day 1 and day 3 then tended towards decrease at day 6 (**Fig. 2F**). Considering these two parameters together, and in light of CG known proliferative behaviour in larval stage (*70*), we propose that adult CG first undergo an endoreplicative phase, between day 1 and day 3, followed by nuclear division between day 3 and day 6. These results suggest that glia development continues beyond the larval stage and proceeds into early adulthood, following a timed pattern of proliferation.

These finding suggested that the decrease in glial volume observed during tumour progression could result from the loss of CG proliferation occurring at day 6. However, quantification of CG nuclei in tumour conditions at day 6 (**Fig. 2E**) revealed a greater loss than expected from proliferation arrest alone, indicating that additional mechanisms contribute to glial loss. Similarly, CG individual nuclear volume in tumour conditions was reduced compared to controls, suggesting a loss of nuclear content (**Fig. 2F**). ltogether, these data show that during tumour progression CG nuclei decrease in volume and in number through mechanisms beyond sole proliferation arrest.

A straightforward explanation would be that glia undergo apoptosis. To test this hypothesis, we took advantage of an apoptotic sensor (*85*), in which the GC3Ai domain (caspase 3-like proteases activity indicator) is fused to an inactive GFP. Upon caspase-3 mediated cleavage, the sensor undergoes a conformational change, allowing GFP fluorescence. We counted all GFP-positive cells as a readout of caspase activation (and further likely apoptosis) during tumour progression compared to control, at days 1, 3 and 6 of adulthood. We found a significant increase in GC3Ai^+^ cells in the tumour context at all timepoints (**Fig. 3. A-B**). We then analysed the spatial distribution of the GC3Ai^+^ cells relative to the tumour and found that the majority of events occurred in close proximity of the tumour (distance of one to two cells), either at the tumour edge or embedded within the tumour, with few GC3Ai^+^ cells located as far as 13 cells away (**Fig. 3C-D**). The more the tumour progressed, the more the distance was mostly shifted to one cell, likely because the expanding tumour occupies most of the tissue, bringing non tumour-related apoptotic events into closer proximity. These findings demonstrate that caspase activation predominantly occurs in cells in close vicinity of the tumour. Co-staining for cell fate markers for glia (Repo) and neuron (ElaV) revealed that most GC3Ai^+^ cells were negative for these markers (**Fig. 3C**), possibly because dying cells downregulate their identity marker before expression of the effector caspase.

**Figure 3.**
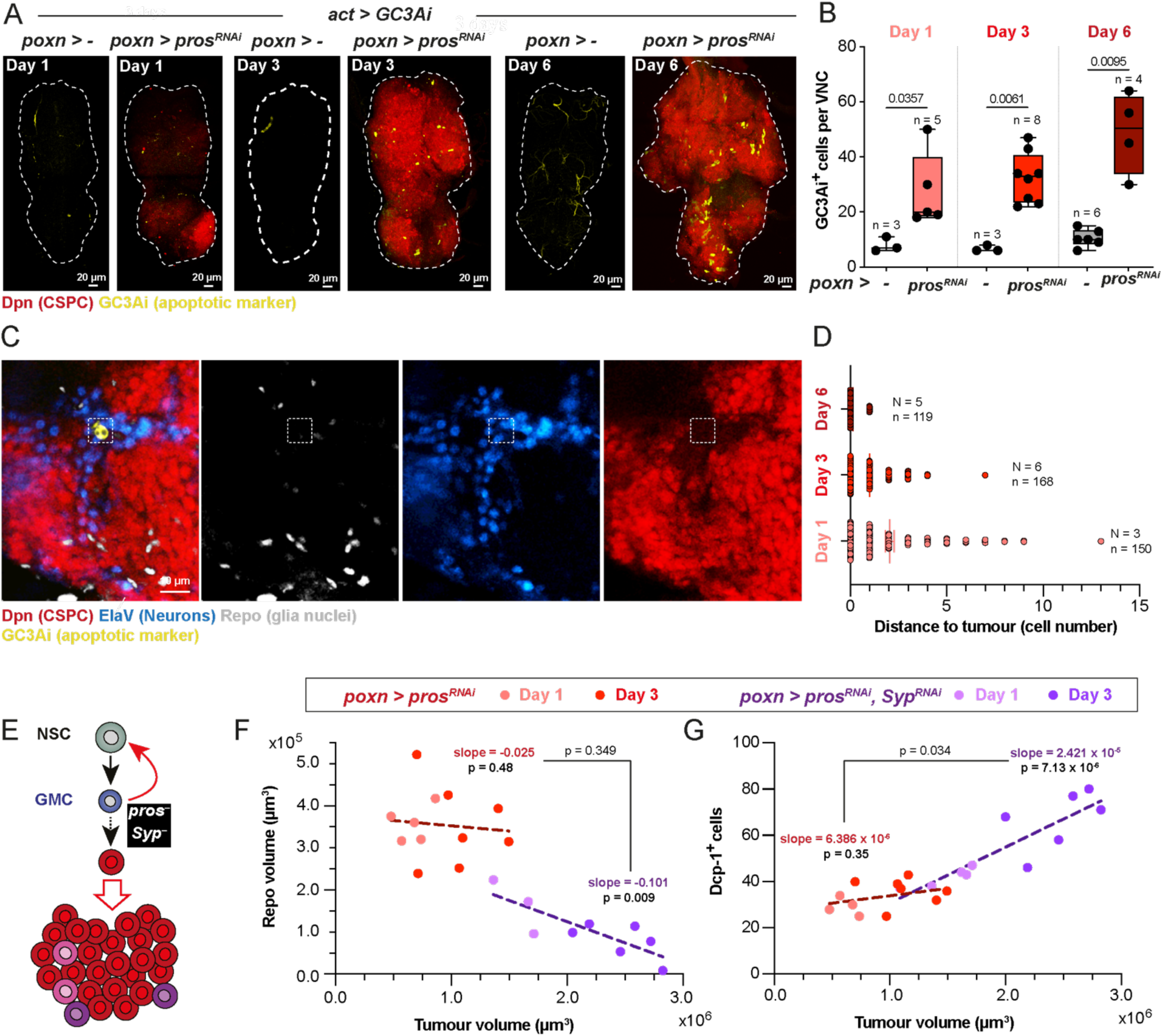
Glia, including the cortex glia, undergo cancer stem cell-driven apoptosis. (A) 3D reconstruction of confocal images showing whole VNCs stained for tumour (Dpn, red) and for a genetic reporter of apoptosis (GC3Ai, yellow) for control (*poxn > -*) and *pros^poxn^* tumour conditions (*poxn > pros^RNAi^*) at days 1, 3 and 6 of adulthood. (B) Box plot of GC3Ai^+^ cells per VNC for control (*poxn > -*) and *pros^poxn^* tumour conditions (*poxn > pros^RNAi^*) at days 1, 3 and 6 of adulthood. n = VNC. Mann-Whitney U tests for each day. (C) Confocal close-up image illustrating the localisation of one apoptotic cell (dashed box), marked by GC3Ai, compared to the tumour edge. This cell has already lost its identity marker, and does not co-stain for glia (Repo) or neuron (ElaV) marker. (D) Dot plot of the number of cells between GC3Ai^+^ cells and the tumour edge at days 1, 3 and 6 of adulthood. Dot = GC3Ai^+^ cells. N = VNC. n = GC3Ai^+^ cells. (E) Schematic of tumour initiation from transformed neural stem cells upon the combined loss of *prospero* (*pros*) and *Syncrip* (*Syp*). These tumours (*Syp^RNAi^, pros^poxn^*) are nearly exclusively composed of CSCs (Imp^+^, Chinmo^+^ and Lin-28^+^). (F) Scatterplot and linear regression analysis of glial volume *versus* tumour volume for *pros^poxn^* tumours (*poxn > pros^RNAi^*) and *Syp^RNAi^, pros^poxn^*(*poxn > pros^RNAi^, Syp^RNAi^)* tumours. Each dot represents a VNC. 12 VNCs were used for *pros^poxn^* tumours and 9 VNCs were used for *Syp^RNAi^, pros^poxn^* tumours to plot the respective linear regression (dotted lines). A linear model was used to calculate the p-values for each linear regression (giving the significance of correlation), and for the difference between the two linear regressions. (G) Scatterplot and linear regression analysis of apoptotic cell counts *versus* tumour volume for *pros^poxn^* tumours and *Syp^RNAi^, pros^poxn^* tumours. Each dot represents a VNC. 11 VNCs were used for *pros^poxn^* tumours and 11 VNCs were used for *Syp^RNAi^, pros^poxn^* tumours to plot the respective linear regression (dotted lines). A linear model was used to calculate the p-values for each linear regression (giving the significance of correlation), and for the difference between the two linear regressions. For box plots, individual values are superimposed. See also Table S1

Given the decrease in glial volume and CG nuclei number and individual volume, together with the increase in caspase activation seen in the TME under tumour condition, we propose that glial cells, including CG cells, undergo apoptosis during tumour progression.

We next sought to assess the relationship between tumour growth and degree of glial loss. To address this, we used the double loss of *pros* and *Syp*, a condition known to produce bigger tumours than *pros* knockdown alone, being nearly exclusively constituted of *Chinmo^+^ Imp^+^ pros^-^* CSCs (*62*) (**Fig. 1A and 3E**). We indeed recapitulated these findings, with *Syp^RNAi^ pros^poxn^* tumours displaying from day 1 more than a twofold increase in tumour volume, followed by an even bigger difference at day 3 (**Fig. S2A-B**). Measuring glial volume (Repo) per VNC showed a more pronounced decrease in *Syp pros^poxn^* tumours compared to *pros^poxn^* tumours, at both timepoints (**Fig. S2C**). In line with this finding, quantifying the number of apoptotic cells using cleaved effector caspase (Dcp-1) staining revealed an increase in *Syp^RNAi^ pros^poxn^* tumours compared to *pros^poxn^* tumours (**Fig. S2D**). Of note, total neuronal volume (ElaV) per VNC did not significantly change in *Syp^RNAi^ pros^poxn^* tumours compared to *pros^poxn^* tumours, although there was a slight tendency towards a decrease (**Fig. S2E**).

These data indicate that glial loss is enhanced in the fast-growing *Syp^RNAi^ pros^poxn^* tumours. We further tested the correlation between tumour and glial volumes in these two tumour conditions. Strikingly, we found no significant correlation between tumour and glial volumes in *pros^poxn^*tumours, whereas these two parameters were negatively correlated in *Syp^RNAi^ pros^poxn^* tumours (**Fig. 3F**). Similarly, when assessing the relationship between tumour volume and the level of apoptosis (as measured by the number of Dcp-1^+^ cells), we found no significant correlation in *pros^poxn^* tumours, but a positive correlation emerged in *Syp^RNAi^ pros^poxn^* tumours (**Fig. 3G**). This shows that glial apoptosis and loss do not occur as direct, linear consequences of tumour growth, and also that they can be shifted to such relationship in *Syp^RNAi^ pros^poxn^*tumours. Notably, *Syp^RNAi^ pros^poxn^* tumours differ in composition from *pros^poxn^* tumours, as they are highly enriched in CSCs (from about 20% to more than 90% of total tumour volume), which in turn causes accelerated tumour growth (*62*). Within this context, our findings suggest that CSC concentration is a critical factor influencing the extent of glial loss, and that CSCs are primary drivers of glial apoptosis.

Altogether, our findings show that CG cells undergo apoptosis in response to tumour growth and propose a cell competition phenomenon whereby tumour cells, and especially CSCs, eliminate their neighbouring glial cells.

### Manipulation of cortex glia apoptosis reveals both its requirement for tumour progression and an intrinsic ability for resistance of the host tissue

We decided to investigate the impact of CG apoptosis on tumour growth. We first assessed the consequence of inducing apoptosis in CG in a precocious manner, at the beginning of adulthood and throughout the whole CG population. We took advantage of the QF binary system to drive the pro-apoptotic *reaper* gene in the CG (*cyp > reaper*) while inducing tumour formation through the GAL4/UAS system, as done previously (*poxn > pros^RNAi^*, *pros^poxn^*tumours) (**Fig. 4A**). The timing of induction was controlled using a heat-inducible cassette (see Methods) to trigger glial apoptosis from adult eclosion (day 0). The increase in cell death events was verified by staining for Dcp-1 at day 6 (**Fig. 4B**). Although it did not reach statistical significance at this timepoint, we observed an upward trend, with some strongly stained VNCs. We then measured the tumour volume overtime by quantifying the Dpn^+^ cells.

**Figure 4.**
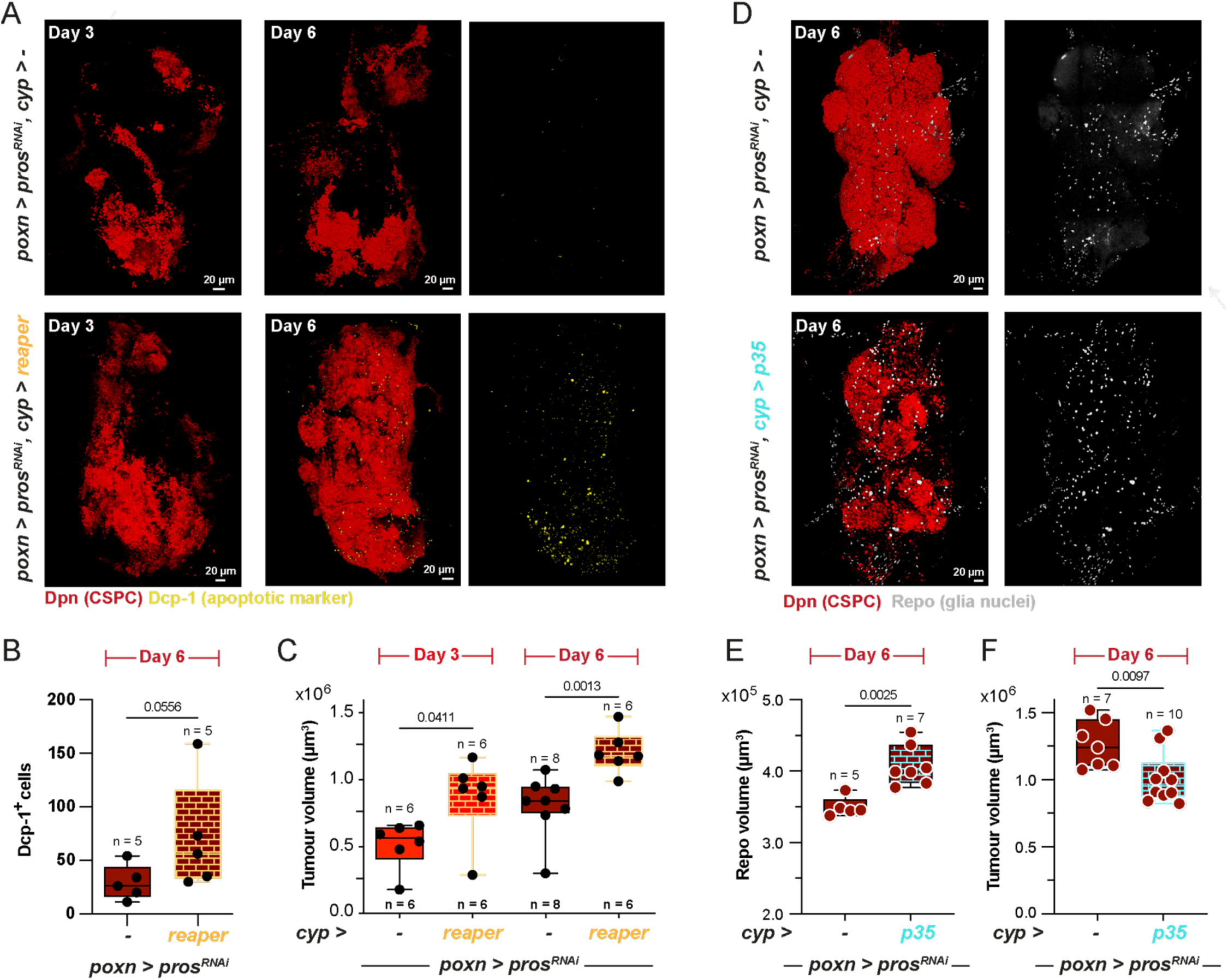
Cortex glia apoptosis promotes tumour growth and its prevention reveals a capacity of resistance from the host tissue. (A) 3D reconstruction of confocal images showing whole VNCs stained for tumour (Dpn, red) and for apoptotic cells (Dcp-1, yellow) for control tumours (*poxn > pros^RNAi^, cyp > -*) and tumours in which apoptosis has been induced from day 0 in all CG cells through the specific expression of the pro-apoptotic gene *reaper* (*poxn > pros^RNAi^, cyp > reaper*) at days 3 and 6 of adulthood. (B) Box plot of apoptotic (Dcp-1^+^) cells numbers per VNC for (*poxn > pros^RNAi^, cyp > -*) and (*poxn > pros^RNAi^, cyp > reaper*) tumour conditions at day 6 of adulthood. n = VNC. Mann-Whitney U test. (C) Box plot of tumour volume for (*poxn > pros^RNAi^, cyp > -*) and (*poxn > pros^RNAi^, cyp > reaper*) tumour conditions at days 3 and 6 of adulthood. n = VNC. Mann-Whitney U tests for each day. (D) 3D reconstruction of confocal images showing whole VNCs stained for tumour (Dpn, red) and for glial cells (Repo, yellow) for control tumours (*poxn > pros^RNAi^, cyp > -*) and tumours in which apoptosis has been blocked from day 0 in all CG cells through the specific expression of the anti-apoptotic exogenous gene *p35* (*poxn > pros^RNAi^, cyp > p35*) at day 6 of adulthood. (E) Box plot of total glial nuclear (Repo) volume for (*poxn > pros^RNAi^, cyp > -*) and (*poxn > pros^RNAi^, cyp > p35*) tumour conditions at day 6 of adulthood. n = VNC. Mann-Whitney U test. (F) Box plot of tumour volume for (*poxn > pros^RNAi^, cyp > -*) and (*poxn > pros^RNAi^, cyp > p35*) tumour conditions at day 6 of adulthood. n = VNC. Mann-Whitney U test. For box plots, individual values are superimposed. See also Table S1.

Remarkably, inducing early and widespread CG apoptosis resulted in a significant increase in tumour volume both at day 3 and day 6 compared to a control tumour condition (**Fig. 4C**).

We then wondered whether preventing apoptosis in glia during tumour progression would have the opposite effect of slowing down tumour growth. To test this, we expressed the anti-apoptotic baculovirus protein, p35 in the CG (*cyp > p35*), from adult eclosion (again relying on the QF system and heat induction, see Methods) and throughout tumour progression (**Fig. 4D**). We first checked whether p35 expression rescued the glial loss witnessed previously. We decided to assess day 6 to ensure that effects of p35 expression could be detected after apoptosis should have been ongoing for some time. We found that glial volume was significantly increased in the tumour condition plus p35 expression in the CG (*poxn > pros^RNAi^*, *cyp > p35*) compared to a control tumour condition (*poxn > pros^RNAi^*, *pros^poxn^* tumours) (**Fig. 4E**). This result confirmed both the effectiveness of p35 induction and our conclusion that CG undergo apoptosis during tumour growth. However, the rescue was only partial when compared to the glial volume measured in non tumour condition (see **Fig. 2C**). It could imply that either p35 expression did not rescue all apoptotic events in the CG or it rescued most but resulted in smaller CG cells (*86*). Alternatively, other glial subtypes may die during tumour growth, or the observed glial loss might also reflect altered proliferation rather than solely apoptosis. We then measured tumour volume under p35 expression in the CG and strikingly we found that at day 6 it was significantly decreased compared to the tumour without p35 expression (**Fig. 4F**).

Altogether these results show that tumour-induced CG apoptosis promotes tumour growth in the host tissue. They also reveal that the CG have the potential to impair tumour progression when they are maintained in the TME.

### Temporal analysis of cortex glia transcriptome reveals a biphasic response to tumour progression

These results highlighted a potential tug-of-war between tumour cells and CG cells, where tumour cells, and in particular CSCs, induce, directly or indirectly, CG cell death, while CG cells exhibit some resistance to tumour progression. To better characterize the behaviour of the CG cells and potentially identify molecular players causing their death or supporting their resistance ability, we determined the transcriptional changes taking place in the CG along tumour growth. We used the Targeted DamID Pol II (TaDa-Pol II) method (*87*, *88*), which allows the tracking of gene expression in specific cell types for given time windows, without having to isolate those cells. This technique uses an enzyme called DNA adenine methyltransferase (Dam), which adds methyl groups to genomic GATC sequences. Fusing Dam with a core subunit of RNA polymerase II (Pol II) results in the methylation of the sequences nearby the polymerase II binding sites, which are mapped to identify paused and active transcription. Controlling its expression by genetic binary systems enables time- and cell population-specific methylation signals. In our case, we placed TaDa-Pol II under the control of the QF system, using a heat-inducible CG-specific driver, while inducing *pros^poxn^* tumours through the GAL4/UAS system (**Fig. S3A**), similarly to the *reaper* and *p35* strategy.

We performed TaDa-Pol II at day 1 and day 6 of adulthood, for both *pros^poxn^* tumour and non-tumour control conditions (**Fig. 5A**). Statistical cross-comparison of the four datasets allowed us to identify gene expression changes in CG between tumour and control conditions over time (**Fig. 5B**; **Dataset S1**). First, we observed a specific response of the CG at day 1 of tumour progression, characterized by a similar number of up- and down-regulated genes. At day 6 however CG display a prominent decrease in gene expression, both compared to the non-tumour condition at day 6 and to the tumour condition at day 1. KEGG pathway enrichment confirmed this biphasic behaviour (**Fig. 5C**; **Dataset S2**). A specific molecular signature of CG cells was revealed at day 1 in tumour context, characterized by an immunity (orange bar) and a lipid metabolism (red bar) components. At day 6 however, most of the downregulated genes belonged to core cellular and metabolic pathways, including oxidative phosphorylation, glycolysis and carbon, tyrosine or glutathione metabolisms (dark blue bar), suggesting a general collapse of essential cell functions. Overall, CG display a biphasic transcriptional response along tumour progression, marked by initial enrichment of specific genes followed by later collapse of core cellular functions.

**Figure 5.**
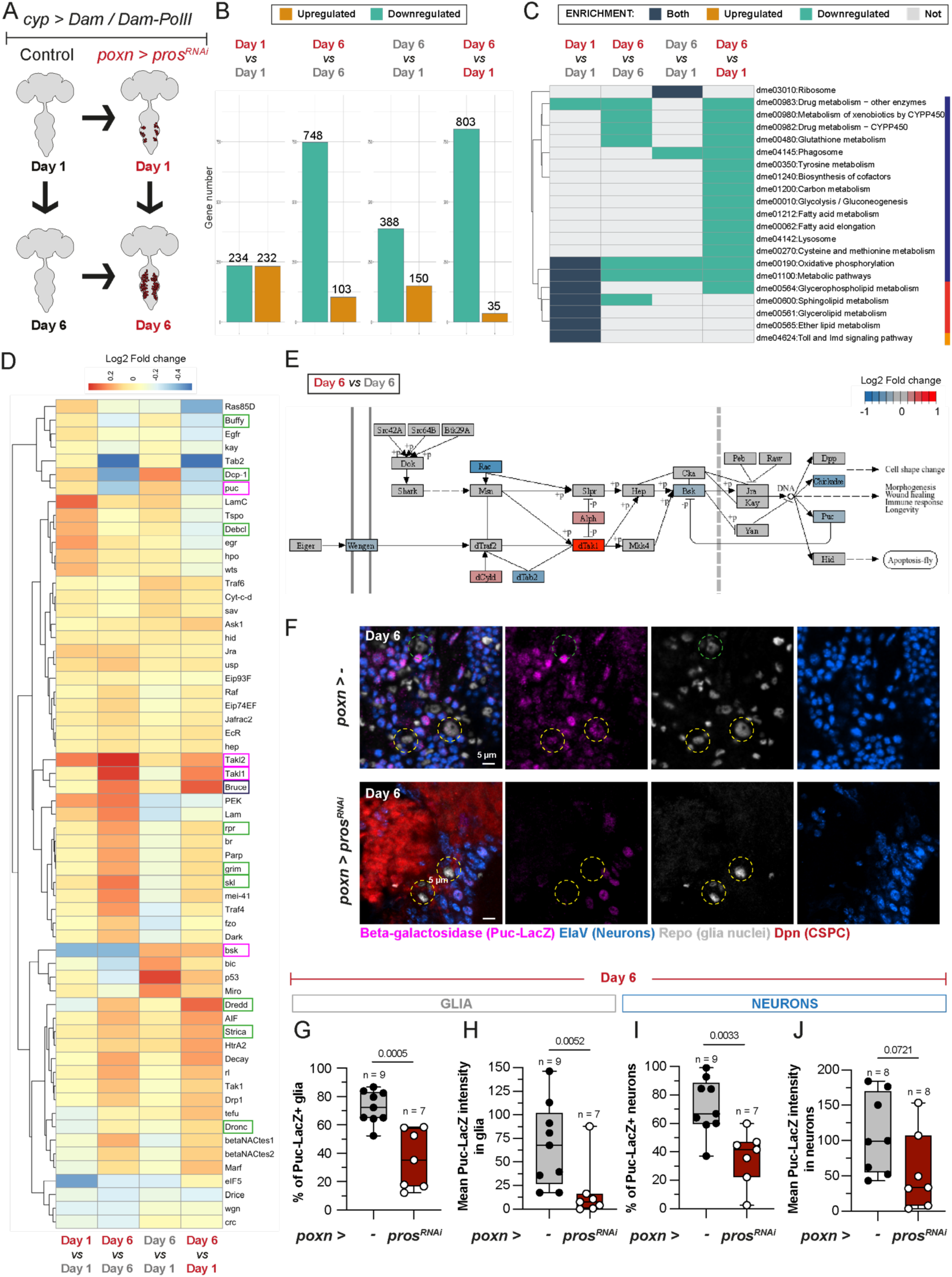
Cortex glia show a biphasic transcriptional response to tumour progression, with an early specific signature shifting to global downregulation of core processes, including the JNK pathway (A) Principle of the comparative Targeted DamID strategy. Either the Dam-PolII fusion, which methylates around PolII binding sites, or the Dam only construct, which accounts for background non-specific methylation, is specifically expressed in the CG under the control of a specific enhancer and using the QF system *(cyp > Dam/Dam-PolII*). Activation of their expression is controlled through heatshock and induced either at day 1 or at day 6, 16 h before dissection. Their combination with *poxn > -* or *poxn > pros^RNAi^* allows the transcriptional profiling of CG cells in control (non-tumour) and tumour conditions respectively, for two timepoints. This produces four datasets in total. Control conditions at days 1 and 6 are coded in black font (*poxn > -, cyp > Dam-PolII* normalized by *poxn > -, cyp > Dam*). Tumour conditions at days 1 and 6 are coded in red font (*poxn > pros^RNAi^, cyp > Dam-PolII* normalized by *poxn > pros^RNAi^, cyp > Dam*). (B) Histograms showing the number of differentially expressed genes between the four different conditions. (C) KEGG pathway enrichment heatmap across the four conditions. The light and dark orange lines indicate the specific early signature (day 1) of CG in tumour conditions, revealing an immune (light orange) and a lipid metabolism (dark orange) components. The dark blue line indicates the overall downregulation in several core metabolic pathways. (D) Differential expression heatmap for genes of the KEGG apoptosis pathway. Fold change scale is in Log2. Green boxes indicate genes of the core apoptotic cascade whose differential expressions support its activation in CG cells upon tumour progression. Dark blue boxes indicate genes of the core apoptotic cascade which are downregulated upon tumour progression. Pink boxes indicate genes of the JNK pathway whose differential expressions support its downregulation in CG cells upon tumour progression. (E) Part of the map of the KEGG *Drosophila* MAPK signalling pathway (Entry dme04013) with differential expression at day 6 between tumour and control conditions overlaid as shades on the gene boxes. Basket (Bsk), the *Drosophila* JNK, and Puckered (Puc), an inhibitor of the JNK pathway whose expression is positively controlled by pathway activation, are both downregulated in tumour condition. (F) Confocal images showing the expression of the *puc-lacZ* reporter (magenta, β-galactosidase staining), whose expression is a readout of JNK pathway activation at day 6 of adulthood in control (non tumour) and *pros^poxn^* tumour conditions. Dashed yellow circles highlight *puc* expression in glial cells. (G) Box plot of the percentage of glial cells positive for *puc-lacZ*/β-galactosidase (Puc^+^ Repo^+^ cells) at day 6 in control (non tumour) and tumour conditions. n = VNC. Mann-Whitney U test. (H) Box plot of the mean intensity of *puc-lacZ* in glial cells positive for *puc-lacZ*/β-galactosidase (Puc^+^ Repo^+^ cells, so excluding Repo^+^ cells with no detected β-galactosidase signal) at day 6 in control (non tumour) and tumour conditions. n = VNC. Mann-Whitney U test. (I) Box plot of the percentage of neurons positive for *puc-lacZ*/β-galactosidase (Puc^+^ ElaV^+^ cells) at day 6 in control (non tumour) and tumour conditions. n = VNC. Unpaired Student t-test. (J) Box plot of the mean intensity of *puc-lacZ* in neurons positive for *puc-lacZ*/β-galactosidase (Puc^+^ ElaV^+^ cells, so excluding ElaV^+^ cells with no detected β-galactosidase signal) at day 6 in control (non tumour) and tumour conditions. n = VNC. Mann-Whitney U test. For box plots, individual values are superimposed. See also Table S1 and Datasets S1 and S2.

In light of our results on CG apoptosis, we first decided to check the expression profiles of members of the apoptotic pathway (**Fig. 5D and S3B**). Analysis across these conditions revealed that several key apoptotic components were upregulated in tumour compared to non-tumour tissues, with different temporal dynamics (at day 1 and/or day 6), fitting our earlier data. This includes core cell death activators *reaper*, *grim* and *sickle* (*skl*), the initiator caspases *dronc*, *dredd* and *strica*, the effector caspase *dcp-1* as well as the pro-apoptotic Bcl2 orthologues, *buffy* and *debcl* (**Fig. 5D**, green boxes). Very few anti-apoptotic factors were upregulated in tumour conditions, such as *bruce* (**Fig. 5D**, dark blue box).

### The JNK pathway is physiologically active in cortex glia and is downregulated during tumour progression

Remarkably, we noticed that several core members of the c-Jun N-terminal kinase (JNK) pathway showed altered expression in tumour conditions (**Fig. 5D**, magenta boxes). This highly conserved MAPK (mitogen-activated protein kinase) pathway is a prominent intracellular player in the response to a range of stress signals and in the regulation of several key cellular processes, including cell proliferation, differentiation, survival, and death. The specific outcomes of JNK activation depend on the stimulus, cell type, and the particular kinases involved. Notably, JNK regulates core pathways playing a crucial role in both cell proliferation and cell death (*89*, *90*), and as such has been since long highly involved in tumourigenesis, in multiple and sometimes conflicating ways (*91–94*). In particular, its known prominent role in cell competition made it a very promising candidate in light of our previous results.

Typically, the activation of MAPK signalling pathways begins when a receptor is activated by a variety of extracellular stimuli (*95*). This triggers the recruitment of adaptor proteins and the activation of small GTP-binding proteins called G-proteins. This induces a cascade of protein, which can consist of up to four successive levels of kinases. For the JNK pathway, JNKKKs (MEKKs, MLKs, DLK, TAK1, ASK1) phosphorylate and activate the JNKKs, which in turn phosphorylate and activate JNKs. Activated JNKs subsequently phosphorylate a diverse array of substrates, including transcription factors (*e.g.,* c-Jun, p53, c-Myc), Bcl-2 family proteins and other kinases and signaling proteins. Analysis of the expression profiles of components of the *Drosophila* JNK pathway showed that the upstream JNKKK dTak1 was upregulated at day 6, whereas JNK itself (*basket, bsk*) was downregulated at day 3 and day 6. Additionally, its negative regulator and transcriptional target *puckered* (*puc*) was downregulated at day 6 during tumour progression (**Fig. 5D**, magenta boxes; **Fig. 5E** and **Fig. S3B**).

*puc* encodes a JNK phosphatase which negatively regulates the activity of the JNK pathway and is itself a transcriptional target of JNK, creating a negative feedback loop (*96*, *97*). As such, *puc* expression has been used as a readout of JNK activation. To assess JNK activity, we relied on the commonly used lacZ enhancer trap line *puc^E69^* (*puc-LacZ*), in which the expression of the β-galactosidase is under the control of *puc* enhancer sequences. This line is also an insertional, homozygous lethal mutant for *puc* (*98*) and therefore, *puc-LacZ* was used in heterozygous. We determined β-Galactosidase (β-Gal) levels in non-tumour (control) and tumour conditions at day 6 of adulthood (**Fig. 5F**), the timepoint when *puc* downregulation is observed in the TaDa-Pol II data (**Fig. 5D**, magenta boxes). Co-staining with Repo to assess β-gal expression in glia revealed that *puc-lacZ* was active in most, though not all, glia cells in control condition. However, in tumour condition, both the proportion of β-Gal^+^ glia and the overall *β-gal* expression levels were sharply reduced (**Fig. 5G-H**). In addition, co-staining with ElaV showed that *β-gal* was expressed in most neurons under control conditions, with higher expression levels than in glia (**Fig. 5I-J**). To our surprise, the proportion of *β-gal^+^*neurons significantly decreased in tumour conditions, accompanied by a reduction in expression level, although less pronounced than in the glia. Of note, no *β-gal* signal was detected in CSPCs suggesting that JNK is not activated in tumour cells (**Fig. 5F**).

Taken together, these data suggest that the JNK pathway is normally active in both glia and neurons, but becomes downregulated in both cell types during tumour growth, with a more pronounced decrease in glia.

### Inhibition of the JNK pathway in CG elicits a dual, temporally dynamic effect on tumour progression and reveals its contribution to host tissue resistance and integrity

JNK pathway activation has been traditionally linked to promoting the elimination of less fit “loser” cells by more competitive “winner” cells in the context of cell competition (*99*, *100*). We thus wondered whether JNK signalling balances resistance and cell competition and how the observed progressive downregulation of JNK in the CG impacts tumour progression. To address this question, we adopted a strategy similar to *reaper* overexpression, blocking JNK activation from the onset of adulthood across the entire CG population. We used the QF system to express a dominant-negative form of Bsk in the CG (*cyp > bsk^DN^*), while simultaneously inducing tumour formation through the GAL4/UAS system (*poxn > pros^RNAi^*, *pros^poxn^* tumours) (**Fig. 6A**). As before, *bsk^DN^* expression was induced by heatshock from adult eclosion (day 0).

**Figure 6.**
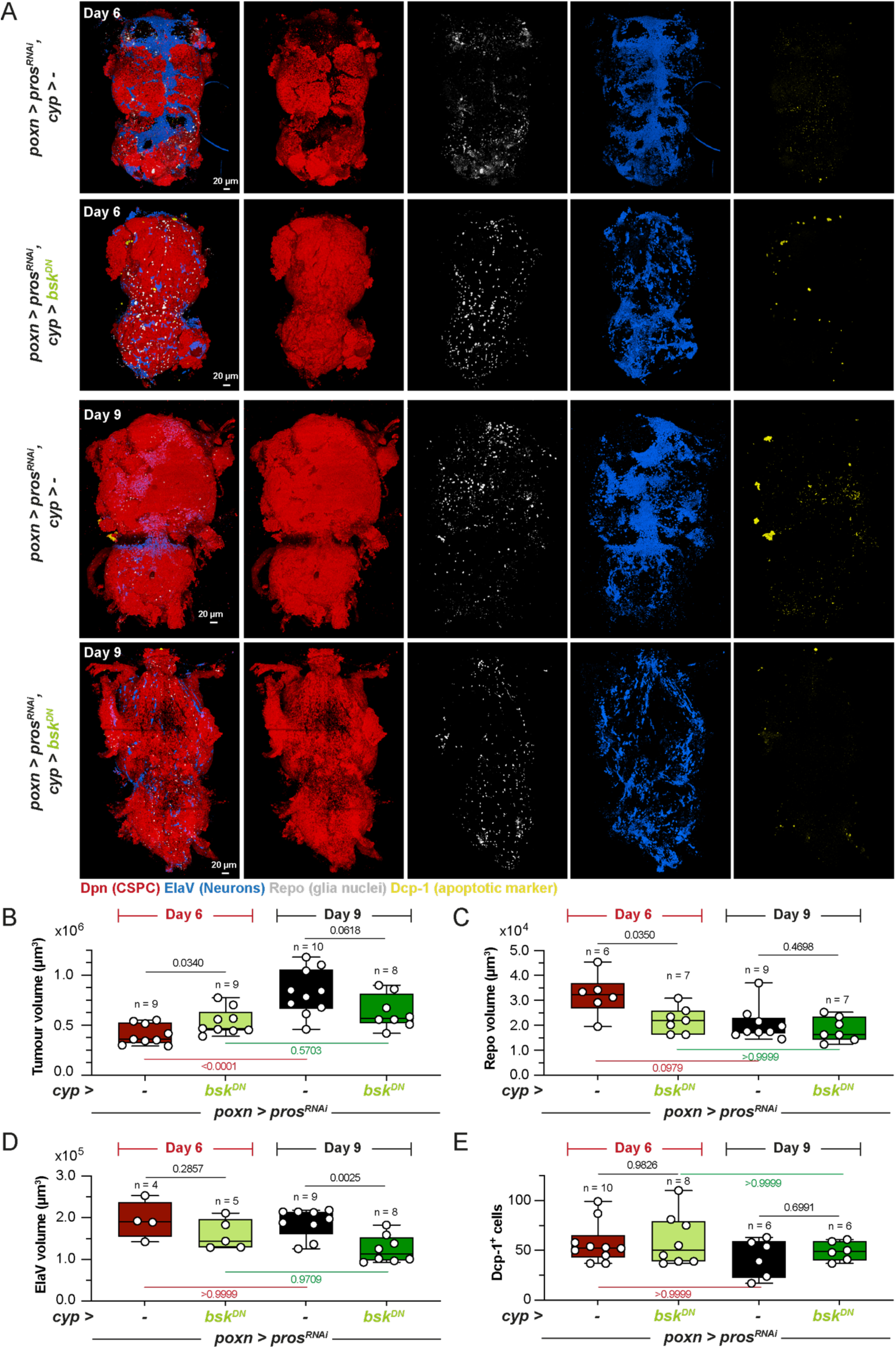
The JNK pathway in CG first hinders then supports tumour growth while protecting neurons from apoptosis. (A) 3D reconstruction of confocal images showing whole VNCs stained for tumour (Dpn, red), glial cells (Repo, gray), neurons (ElaV, blue) and apoptotic cells (Dcp-1, yellow) for control tumours (*poxn > pros^RNAi^, cyp > -*) and tumours in which a dominant negative form of Bsk has been induced from day 0 in all CG cells (*poxn > pros^RNAi^, cyp > bsk^DN^*) at days 6 and 9 of adulthood. (B) Box plot of tumour (Dpn) volume for (*poxn > pros^RNAi^, cyp > -*) and (*poxn > pros^RNAi^, cyp > bsk^DN^*) at days 6 and 9 of adulthood. n = VNC. One-way ANOVA with with Tukey’s multiple comparisons test between days. Student’s t-tests between control and *bsk^DN^* conditions for each day. One outlier was removed for (*poxn > pros^RNAi^, cyp > bsk^DN^*) at day 6. (C) Box plot of glia (Repo) volume per VNC for (*poxn > pros^RNAi^, cyp > -*) and (*poxn > pros^RNAi^, cyp > bsk^DN^*) at days 6 and 9 of adulthood. n = VNC. Kruskal-Wallis with with Dunn’s multiple comparisons test between days. Mann-Whitney U tests between control and *bsk^DN^* conditions for each day. (D) Box plot of neuron (ElaV) volume per VNC for (*poxn > pros^RNAi^, cyp > -*) and (*poxn > pros^RNAi^, cyp > bsk^DN^*) at days 6 and 9 of adulthood. n = VNC. Kruskal Wallis with Dunn’s multiple comparisons test between days. Mann-Whitney U tests between control and *bsk^DN^* conditions for each day. (E) Box plot of apoptotic cell (Dcp-1) numbers per VNC for (*poxn > pros^RNAi^, cyp > -*) and (*poxn > pros^RNAi^, cyp > bsk^DN^*) at days 6 and 9 of adulthood. n = VNC. Kruskal Wallis with with Dunn’s multiple comparisons test between days. Mann-Whitney U tests between control and *bsk^DN^*conditions for each day. For box plots, individual values are superimposed. See also Tables S1 and S2.

The analysis at day 6 of tumour volume (Dpn^+^ cells) when *bsk^DN^* was expressed in the CG (*poxn > pros^RNAi^*, *cyp > bsk^DN^*) revealed a mild yet significant increase compared to a control tumour condition (*poxn > pros^RNAi^*, *pros^poxn^* tumours) (**Fig. 6B**). This was accompanied by a significant decrease in both glial volume (**Fig. 6C**) and nuclei number (**Fig. S4A**). Glial individual nuclear volume was actually increased (**Fig. S4B**), reflecting a shift in the glial population, with CG being depleted and other subtypes possessing large polyploid nuclei, such as subperineurial glia, becoming relatively enriched. No significant changes were detected in neuronal volume (**Fig. 6D**) or cell apoptosis (number of Dcp-1+ cells, **Fig. 6E**), despite a slight trend towards decreased neuronal volume. These results indicate that active JNK signalling in CG does not induce apoptosis nor cell elimination, but instead plays an anti-tumour role, constraining its growth.

Based on these findings, we next evaluated the impact of JNK function at a later time point, day 9 of adulthood, anticipating a stronger effect on the various cell populations. Suprisingly, we found a clear, significant decrease in tumour volume when *bsk^DN^* was expressed in the CG, opposite to the increase observed at day 6 (**Fig. 6B**). We calculated that the mean growth rate of control tumour between day 6 and day 9 was close to 2, while tumours expressing *bsk^DN^* in the CG had a reduced mean growth rate of 1.2 (see **Table S2**). These data indicate that JNK signalling in CG becomes beneficial to tumour growth at later stages, and that its sustained downregulation ultimately hinders tumour growth. This pro-tumour role was not accompanied by any change in glial volume or apoptotic cell counts compared to control tumours (**Fig. 6C, E**), suggesting that it is independent of CG elmination. However, we recorded a significant decrease in neuronal volume at day 9 when *bsk^DN^* was expressed in the CG (**Fig. 6D**). Importantly, neuronal volume was not decreased in control tumours compared to non-tumour conditions at day 9, showing that at this stage neurons remain protected from tumour-induced loss when JNK is active (**Fig. S4C-D**). Thus JNK signalling in CG provides a non-cell autonomous protection to neurons during tumour progression.

We then wondered whether this neuroprotective role was specific to tumour conditions, or also present under physiological, non tumour conditions. We induced *bsk^DN^* in CG from day 0 of adulthood using the TARGET system, in which the onset of GAL4 activity is controlled through temperature shift by a thermosensitive version of its inhibitor GAL80 (*101*) (**Fig. S5A**). We measured the volume, nuclei number and average nuclear volume of glial cells (Repo^+^) per VNC, finding no significant differences between control and *bsk^DN^*except at day 9 (at 29°C), when *bsk^DN^* expression led to more glial nuclei of smaller volume (**Fig. S5B-D**). Neuronal volume also remained largely unchanged between control and *bsk^DN^*, although a slight, non-significant decrease was observed at day 9 (**Fig. S5E**). However, expressing *bsk^DN^* in the CG resulted in higher numbers of apoptotic Dcp-1^+^ cells in the VNC at all time points examined (**Fig. S5F**). These results suggest that physiological JNK signalling may exert a mild neuroprotective effect by limiting apoptosis, consistent with our observations under tumour conditions.

Together, these data reveal that during adulthood, JNK signalling is active in glial cells where it has a neuroprotective function and provides partial resistance to tumour growth. However, during tumourigenesis, JNK signalling is silenced in glia surrounding the tumour, leading to increased tumour growth and consequently reduced glial numbers. While JNK silencing in CG during tumourigenesis initially facilitates tumour growth, in the long term it contributes to a global collapse of the nervous system along with restricting tumour progression.

## DISCUSSION

Using a *Drosophila* CSC-driven tumour model, this study explores how host cells within the TME influence tumour progression. We first found that the cortex glia (CG), a major glial subtype in direct contact with the tumour, remodel their membranes and infiltrate the tumour mass during tumour growth. However, this infiltration declines over time, concurrent with CG cells undergoing apoptosis and subsequent elimination. Strikingly, CG cell death plays a pivotal role in tumour progression, as accelerating or preventing it, increases or decreases tumour growth respectively. This finding reveals that CG cells possess an intrinsic capacity for resistance, which tumour cells ultimately overcome via CSC-driven cell competition, allowing tumour expansion. These two tumour-host interaction phases are reflected by the CG transcriptome over time. Initially, a specific CG signature is present, but it later shifts to a general downregulation of core cellular functions and metabolic pathways, including the JNK pathway. We found that the role of JNK downregulation in tumour progression is also two-faced. Initially, active JNK signalling confers resistance to tumour growth, and its downregulation is thus beneficial to the tumour. However sustained downregulation ultimately appears detrimental to the tumour, restricting its growth. Importantly, his latter phase is accompanied by neuronal loss, revealing a neuroprotective role of JNK signalling in the CG. Thus, our study highlights interlinked biphasic behaviours at both host tissue and tumour levels, which shape tumour growth dynamics. The host tissue possesses resistance mechanisms which are eventually overcome by sustained CSC-driven competition, ultimately leading to collapse of host tissue and its supportive functions, and curbing tumour growth itself (**Fig. 7A-C**).

**Figure 7.**
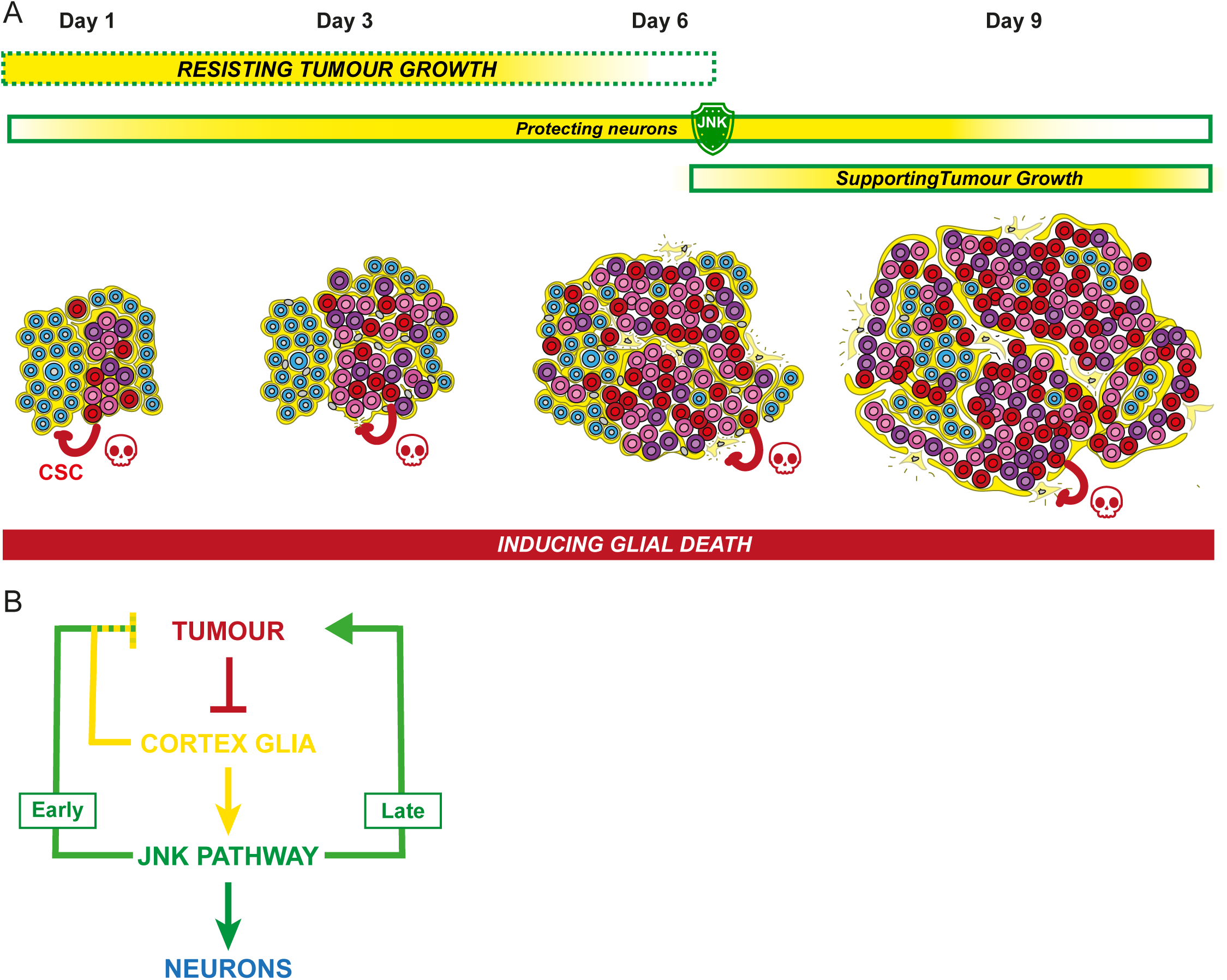
Dynamics of interplay between cortex glia cells and tumour progression (A) *pros^poxn^* tumours, driven by CSCs (red), and whose bulk is made of CPCs (pink and purple) grow extensively in a TME made of CG cells (yellow) and neurons (blue) between day 1 and day 9 of adulthood. Tumour growth triggers the remodelling of CG membranes, which infiltrate within each tumour mass while keeping the original enwrapping at its boundary. In addition, CSC-driven tumour progression induces apoptosis of the CG cells (red arrows), what decreases the density of CG infiltration. This creates a dynamic balance: CG death favours tumour growth, while CG presence provides resistance to such growth. Early resistance relies in part on intrinsic, pre-existing JNK signalling (green coded) in the CG, a pathway which also protects neurons from death. Yet other mechanisms, regulated by the JNK pathway, also exist in the CG to play an opposite supportive role for tumour growth in later phases - possibly by preventing the collapse of the host tissue and its diverse support functions. (B) Tumour growth leads to CG apoptosis, a cell loss which results in the overall decrease of JNK signalling normally active in the CG cells. Initially, intrinsic JNK activity provide some resistance to tumour growth, and its downregulation further favours tumour growth, setting up an amplifying loop (Early). However, over time JNK signalling becomes essential for tumour support and its loss in CG is detrimental to tumour progression, which starts to stall (Late). The latter phase coincides with neuronal loss, supporting a neuroprotective role of the JNK pathway in CG.

### Growth of tumour clones

The tumour, observed in the adult stage as a whole, is actually the result and sum of the growth of six individual tumours from larval stage. Using a multicolour clonal analysis, we found that these tumours do not seem to intermingle but remain well-delineated, with CG membrane separating them, at least during the time of our analysis (**Fig. 1J-K** and **Fig. S1B-D**). In larval stage, normal NSC lineages rely on differential adhesion to prevent merging with neighbouring lineages, independent of CG wrapping, and individual *pros* tumours similarly remain separated (*78*). In our model, it would be interesting to see whether this clonal organisation is indeed intrinsic and controlled by adhesion complexes, or relies on the maintenance of the CG barrier between clones. Future studies on the adhesive molecular signature of these tumours are needed to better understand the inter-clonal organisation, and could provide insight into why and how CG membranes can infiltrate the tumour at this stage.

### Cancer stem cell-driven cell competition

Previous studies on epithelial tumours in the *Drosophila* gut and in mouse intestinal organoids have shown that tumour mutant clones kill their healthy counterparts, a cell competition-based phenomenon essential for tumour growth (*42*, *44*). Here, we find that *pros^poxn^* tumours progressively induce apoptosis and elimination of CG cells, and that, in turn, glial death promotes tumour growth. Cell competition-induced death within the TME therefore could be a fundamental process driving tumour growth across various cancers –including CNS tumours, where this mechanism had not been previously identified and where host cells are non-epithelial and of different origin.

Yet, molecular mechanisms appear to differ. In gut epithelial tumours, the JNK pathway is active both in tumour cells and surrounding cells, where it is required for their growth and their elimination, respectively (*42*). Several other epithelial contexts of cell competition, with genetically-induced mosaic populations of either less fit/loser (*eg*, *minute* (*40*), *scrib* mutants (*102*, *103*)) or super-competitor/winner (*eg*, *myc* (*104*, *105*)) cells *versus* wild-type cells (thus respectively behaving as winner or loser), have shown that JNK pathway activation in loser cells is key for their engagement into apoptotic processes and elimination (reviewed in (*39*, *99*, *100*, *106*, *107*)). When wild-type cells act as winners, JNK is also active in these cells, mediating the engulfment of loser cells. Finally, models of compensatory proliferation following apoptosis (also known as apoptosis-induced proliferation) have involved JNK pathway activation in the apoptotic and/or responsive cells, this time downstream of apoptosis (*100*, *106*, *108*).

In the CNS, JNK is not active in tumour cells (**Fig. 5F**). Moreover, JNK signalling is constitutively on in CG cells, rather becomes downregulated during tumour progression (**Fig. 5**), and does not contribute to CG elimination, as its inhibition fails to rescue glial loss (**Fig. 6C**). This highlights a role of JNK beyond its established functions in cell competition. How tumour cells induce apoptosis in CG cells remain to be elucidated. Potential mechanisms include direct cell-cell interactions through transmembrane partners, paracrine short-range diffusive signals and mechanical stress. Our finding that the concentration in CSCs, rather than overall tumour volume *per se*, correlates with the level of glial loss (**Fig. 3F-G**) is striking and suggest that CSCs are essential and likely a key source of (possibly paracrine) apoptotic signals.

### Glial cells and JNK signalling spearhead host tissue resistance to tumour growth

Although competition from the tumour ultimately eliminates and overpowers CG cells, these cells exhibit a certain potential to hinder tumour progression. This resistance to tumour growth (and not to death) is demonstrated by the effects of either removing or preserving CG cells in the TME (**Fig. 4A-F**). This exciting finding suggests that targeting the host tissue, and, perhaps counterintuitively, blocking cell death, could represent an interesting angle for therapeutic approaches. A prerequisite would be to identify the mechanisms behind such resistance, which may be multifactorial and not mutually exclusive, including mechanical- and signalling-driven processes. One intrinsic, already present mechanism seems to involve the JNK pathway. JNK signalling is normally active and present in CG, where it where it exerts an anti-tumour effect, at least during the initial stages (**Fig. 6A-B**). This anti-tumour role does not seem to involve apoptosis (**Fig. 6E**), and other known functions of JNK signalling, such as regulation of proliferation and cytoskeleton organisation, require further investigation. Beyond already pre-existing resistance mechanisms, CG could actively set up additional defense or attack systems in response to tumour growth. The existence of a distinct transcriptomic signature early during tumour progression (day 1) (**Fig. 5B-C**) may provide insight into these mechanisms. This signature is characterized by the upregulation of immunity-related genes, including antimicrobial peptides, and genes involved in lipid metabolism. It has been shown that the *Drosophila* immune system reacts to and kill epithelial tumour cells with the release of antimicrobial peptides (*109*), suggesting that their upregulation in the CG could contribute to targeting the tumour. Moreover, activation of conserved immune pathways across diverse competition models has been identified as a driver of loser cells apoptosis (*110*). However the functional relevance of this genetic signature for CG resistance will have to be disentangled from other opposite roles of the CG in tumour support. This is a general interrogation both for pre-existing or set up cellular and molecular processes, as we uncovered with the biphasic role of the JNK pathway.

### Late JNK signalling: turning pro-tumour or avoiding whole system’s collapse?

Unexpectedly, we found that JNK downregulation led to a dramatic slowdown, nearly to a halt, of tumour growth at a later phase (**Fig. 6A-B** and **Table S2**), showing JNK switches to a pro-tumour role. Theis finding highlights the dual role of the TME in regulating tumour progression. JNK in the CG could help tumour growth in many ways, by trophic support, architectural remodeling, or signaling mechanisms. Notably, we observed an infiltration of the tumour clones by CG membranes, something we do not see for non-transformed lineages or very small tumours (**Fig. 1D, H, K** and (*78*)). CSCs may acquire some properties (signals and/or loss of inter-adhesions) which attract or/and are permissive for CG to infiltrate; in turn such infiltration could sustain tumour growth by locally providing important factors. Interestingly, in a glioma model (*111*) where CG cells themselves undergo transformation, JNK activation promotes CG membrane remodeling and the formation of an infiltrating interconnected network which support tumour growth while causing neurodegeneration (*112*). It would be interesting to test whether JNK signalling regulates CG membrane network in our model.

Why the loss of JNK signalling is ultimately detrimental to tumour growth, as such a late stage, is left to be investigated. An intriguing observation is that such effect is concomitant to the loss of neurons, whose survival appears to depend on JNK signalling in the CG (**Fig. 6A, D**). Of note, JNK signalling in glia promotes neuronal phagocytosis and clearance during *Drosophila* development (*113*, *114*), highlighting complex neuron-glia dynamics. In adult, JNK neuroprotective role also takes place in normal, non-tumour condition, although to a lesser extent (**Fig. S5E, F**). A compelling hypothesis is that neuronal loss is causative of tumour slowdown, rather than JNK directly promoting tumour progression. Recent studies have indeed connected the presence and activity of neurons to the promotion of tumour growth, emphasizing the cooption of normal neuron-glia interaction in these processes (reviewed in (*27*, *28*)). Our model could fit in this picture, with tumour growth dependent on neurons.

Notably, neurons in control tumours do not seem to die until day 9, suggesting that they are either shielded from tumour growth longer than glial cells or that neuronal loss is too minimal to detect. Yet, the slight trend towards neuronal volume decrease for *Syp^RNAi^pros^poxn^*tumours (**Fig. S2E**), where tumours grow and glial cells die much faster, may suggest that neurons could eventually be lost as well. It would be interesting to check neuronal volume and apoptosis beyond day 9 in *pros^poxn^*tumours to determine whether neuronal loss eventually occurs and coincides with host death. In *pros^poxn^* tumours, the death of CG cells reduces JNK signalling at the tissue level, and surviving glial cells exhibit lower JNK activity, as indicated by reduced *puc* expression in TaDa-Pol II profiling and reporter assays (**Fig. 5D, F-H**) –what would dramatically decrease JNK signalling on the long run and affect neurons.Thus, the level of JNK signalling may be a key mechanism deciding on neuronal fate under tumour condition, rather than sheer tumour growth alone. Curiously, JNK levels are also lowered in neurons, albeit less than in glial cells (**Fig. 5F, I-J**). It remains unclear whether JNK downregulation in one cell type influences the other, or what ultimately triggers cell death. Nevertheless, the role of JNK signalling in CG for neuronal survival is clear (**Fig. 6D**), independently of potential relays. Ultimately, neuronal fate may determine the overall trajectory of tumour progression

By analysing cell types, cellular processes, gene expression profiles and pathway functions, our study converges on the critical importance of timing in tumour growth, specifically highlighting multiple biphasic behaviours. Within the tumour microenvironment, glial cells transition from an initial resistance and active response phase to eventual collapse, while the JNK pathway exerts a dual, progressive influence on both tumour and neurons. The complex, dynamic and sometimes antagonstic functions of the different cellular and molecular players in the TME interact and compete with the CSC-driven tumour. The timing of these evolving interactions will shape tumour growth and outcome. This central concept, together with the presence of intrinsic resistance mechanisms in the host tissue, highlights the importance of early intervention in tumour development. Enhancing host tissue defense mechanisms emerges as a therapeutic alternative to directly targeting the tumour.

## METHODS

### Fly lines and husbandry

*Drosophila melanogaster* lines were kept and raised in vials containing standard cornmeal food medium. Two copies of each stock were maintained at 18°C and flipped into fresh food vials every 4-6 weeks until their use in crosses and experimental procedures. Working stocks were kept at 25°C.

Parental lines were crossed in vials and allowed to lay eggs for 2 days at 25°C, then flipped in another vial. Larvae and pupae from each egg laying were left to develop in these vials at 25°C, unless specified in Table S1. Day 0 of adulthood was defined as at the day of eclosion from the pupal case for each adult.

18°C was used to keep the thermosensitive allele of GAL80 (GAL80^ts^), a repressor of GAL4, active (to switch off GAL4 expression), while 29°C was used for its inactivation (to switch on GAL4 expression). Temperature switch was performed at eclosion (day 0).

For induction of QUAS-reaper, QUAS-p35, QUAS-bskDN and QUAS-puc, adult virgin females at eclosion (day 0), were heat shocked for 4h at 37 °C and aged to the desired day before dissection.

Detailed genotypes, crosses and culture regimens are listed in Table S1. All the experiments were performed only on female flies.

The lines used in this study are listed below:

| Strains | Source | Stock number/Reference |
| --- | --- | --- |
| *w<sup>1118</sup>* | BDSC | 5905 |
| *yw, hs-FLP* | Andrea Brand lab |  |
| *Nervana2::GFP (Nrv2::GFP)* | BDSC | 6828 |
| *puc<sup>E69</sup> (puc-lacZ)* | Stéphane Noselli lab | (98) |
| *cyp4g15(cyp)-myr::tdTomato* | Spéder lab | (70) |
| *tubulin-GAL80<sup>thermosensitive(ts)</sup>* | BDSC | 65406 |
| *cyp4g15(cyp)-GAL4* | BDSC | 39103 |
| *poxn-GAL4* | BDSC | 66685 |
| *actin-LexA* | BDSC | 62567 |
| *cyp4g15(cyp)-QF2* | Pauline Spéder lab | (70) |
| *cyp4g15-FRT-STOP-FRT-QF2* | Pauline Spéder lab | This study |
| *UAS-dicer* | BDSC | 24650 |
| *UAS-bsk<sup>DN</sup>* | BDSC | 6409 |
| *UAS-prospéro RNAi* | VDRC | 101477 |
| *UAS-Syncrip-RNAi* | VDRC | 33011 |
| *UAS-raeppli-NLS 53D* | BDSC | 55087 |
| *LexAOp-GC3Ai* | Romain Levayer lab | (85, 115) |
| *QUAS-His2Av::mCherry* | BDSC | 93911 |
| *QUAS-Dam* | Pauline Spéder lab | This study |
| *QUAS-Dam-PolII* | Pauline Spéder lab | This study |
| *QUAS-bsk<sup>DN</sup>* | Hermann Steller lab |  |
| *QUAS-p35* | Hermann Steller lab | (116) |
| *QUAS-reaper* | Hermann Steller lab | (116) |

### DNA cloning and *Drosophila* transgenics

*cyp4g15-FRT-STOP-FRT-QF2* was generated in the same fashion as *cyp4g15-FRT-STOP-FRT-LexA* (*70*). Part of the cyp4g15 enhancer (GMR55B12, Flybase ID FBsf0000165617) was amplified from genomic DNA extracted from cyp4g15-GAL4 adult flies, with a minimal Drosophila synthetic core promoter [DSCP93] fused in C-terminal. A FRT STOP cassette was amplified from an *UAS-FRT.STOP-Bxb1* plasmid (gift from MK. Mazouni) and the QF sequence was amplified from the entry vector pENTR L5-QF2-L2 (gift from S. Stowers / M. Landgraf; Addgene 32305). The two amplicons were joined by overlapping PCRs. This *FRT-STOP-FRT-QF2* amplicon was inserted together with the *cyp4g15^DSCP^* enhancer in the destination vector pDESThaw sv40 (gift from S. Stowers) using the Multisite gateway system (*117*) to generate a *cyp4g15^DSCP^-FRT-STOP-FRT-QF2* construct. The construct was integrated into the fly genome at either an attP2 or attP40 docking site through PhiC31 integrase-mediated transgenesis (BestGene). Several independent transgenic lines were generated and tested, and one was kept.

QUAS-Dam and QUAS-Dam::PolII were created by amplifying the Dam and Dam::PolII sequences respectively from the pUASt-attB-LT3-Dam and pUASt-attB-LT3-Dam::PolII plasmid (*87*). Each amplicon was inserted into the entry vector pDONR P5-L2 to create entry vectors pENTR-L5-Dam-L2 and pENTR-L5-Dam::PolII-L2. Each of these pENTR was combined with the entry vector pENTR-L1-10XQUAS-R5 and the destination vector pDESThaw sv40 (both gifts from S. Stowers) using the Multisite gateway system (*117*) to generate the QUAS-Dam and QUAS-Dam::PolII constructs. The constructs were integrated into the fly genome at either an attP40 docking site through PhiC31 integrase-mediated transgenesis (BestGene). Several independent transgenic lines were generated and tested, and one was kept.

### Transcriptional profiling

We used Targeted DamID (*87*, *88*, *118*) to profile transcription in cortex glia at different timepoints, during tumour progression. *QUAS-Dam* and Q*UAS-Dam::Pol II* flies were crossed to *tubulin-GAL80^ts^*; *cyp4g15-GAL4* flies. Induction was performed through a 4 h heat shock at 37°C at day 1 and day 6 of adulthood. The flies were dissected after 14 h of induction and the VNCs were stored at –80°C in 50ul of PBS, until samples were ready to be prepared for the DNA extraction. Processing of genomic DNA were performed as previously described (*88*). Two biological replicates were performed for each timepoint. For each condition, DamID-seq samples were processed via the damid_seq pipeline v1.5.3 (*118*), that automatically handles sequence alignment, read extension, binned counts, normalization, pseudocount addition and background reduction (Pol-II / Dam only). Average Pol-II occupancy and FDR were calculated for each gene of Drosophilia melanogaster genome (DmR6 release) with the polii.gene.call script (*88*, *119*). Differentially expressed genes were identified via the NOIseq R package (*120*): RNA pol II gene occupancy scores were scaled, and inverse log values used as input to NOIseq with parameters of upper quantile normalization and biological replicates. DEGs were called with a q-value of 0.90. The functional enrichment of up and down regulated genes was calculated with the DAVID analysis tool (*121*) using Gene Ontology, KEGG and Reactome functional gene sets and Drosophila melanogaster annotations as background. Annotation categories were considered as enriched when at least 2 genes were involved in and passing the EASE score threshold of 0.1. Treemap plots and heatmaps were generated using R (v4.1.3) (*122*) and the treemap (v2.4-3) (*123*) and pheatmap (v1.0.12) (*124*) packages. KEGG maps have been designed with Pathview R package (*125*).

### Fixed tissue immunostaining

Adult CNSs were fixed for 30 minutes in 4% methanol-free formaldehyde (ThermoFisher 28908) while rocking at room temperature. They were washed in PBS once and permeabilized in PBS containing 1% Triton (Sigma T9284) three times for 10 minutes each. CNSs were incubated with blocking solution (PBS-Triton 1%, 5% Bovine Serum Albumin (Sigma A3608), 2% Normal Goat Serum (Abcam ab7481) for 1 hour at room temperature while rocking. Primary antibody dilution was prepared in blocking solution and CNSs were incubated at 4°C for 48 hours, then washed three times with PBS-Triton 1% while rocking. Secondary antibodies were diluted in blocking solution and CNSs were incubated at room temperature for 2 to 3 h. CNSs were washed three times with PBS while rocking and kept in PBS at 4°C until mounted.

Homemade mowiol mounting medium (Sigma 81381) was used to mount the CNSs between borosilicate glass slides (number 1.5; VWR International) and coverslips. Primary antibodies used were: guinea pig anti-Dpn (1:1000,(*70*)), rat anti-ElaV (1:100, 7E8A10-c, DSHB), mouse anti-Repo (1:100 (DSHB, 8D12-c), rabbit anti-Dcp-1 (1:100, Cell Signaling 9578), rabbit anti-β-galactosidase (1:200, ThermoFisher Scientific A-11132). Fluorescently-conjugated secondary antibodies Alexa Fluor 405, Alexa Fluor 488, Alexa Fluor 546 and Alexa Fluor 633 (ThermoFisher Scientific) were used at a 1:200 dilution. DAPI (4′,6-diamidino-2-phenylindole, ThermoFisher Scientific 62247) was used to counterstain the nuclei.

### Image acquisition and processing

Confocal images were acquired using a laser scanning confocal microscope (Zeiss LSM 880, Zen software (2012 S4)) with a Plan-Apochromat 40x/1.3 Oil objective. All VNCs were imaged as z-stacks with an optimal distance between each slice of 1 μm and using tiles, with 5-10 % overlap. Images were analysed and processed using Fiji (Schindelin, J. 2012), Volocity (6.3 Quorum technologies), and Imaris (version 10.2.0). Denoising was used for some images using the Remove noise function (Fine filter) in Volocity. Images were assembled using Adobe Illustrator 29.6.1.

### Quantification of tumour, glial and neuronal volumes

Dpn, Repo, ElaV and His::mCherry (under the control of *cy4g15* enhancer sequences, *cyp > His::mCherry*) signals were used to measure nuclear volumes for tumour, glia, neurons and CG cells respectively. For Repo and His::mCherry signals, in which the nuclei are better separated in space, the nuclear volume for individual cell was also calculated.

Two different protocols were used.

For Fig. 1C, 2B-C, S2, 3F-G and 4, nuclei volume quantification was achieved with a Python script deposited on Zenodo (10.5281/zenodo.16733817) together with its operating manual.

For Fig. 6, S4 and S5, raw confocal Zeiss files (.czi) were opened in Imaris (version 10.2.0). Once converted to an Imaris *.ims file, nuclei segmentation was performed by applying the Surface Wizard tool to the desired cell channels. The following settings were used for all the images: 0.415 μm for smooth surface details; background subtraction and the diameter of largest sphere was 1.56 μm; enabled split touching object and morphological split; seed points diameter 2.08 μm. The processed images were visually examined alongside the original channels to ensure correct identification of cell types. Statistics for individual images were exported in either Microsoft Excel or CSV format. The data were then imported into GraphPad Prism for quantitative/statistical analysis.

### Quantification of Puc-lacZ signal

Image analysis to assess puc expression level in glia and neurons was performed on Fiji to quantify volumetric and fluorescence intensity data from 3D image stacks, following four steps.

1. Image Selection and Preprocessing. Individual channels (Puc-lacZ and either Repo or ElaV) were duplicated two times. Thresholding was determined manually and applied to each channel, resulting in image pairs of the original channel image and the threshold binary mask for *puc* expression and the cell population of interest (for exemple: Puc threshold Mask 1 and its corresponding fluorescence image with Elav threshold Mask 2 and its corresponding fluorescence image). These masks are used to define regions of interest (ROIs) for subsequent analysis.
2. Mask Combination and Volume Calculation. An intersection of the two selected masks was generated using the “AND create stack” operation in Fiji (Image Calculator tool), resulting in a combined mask (“Puc and Repo or ElaV Mask”). The individual and combined volumes of Puc-lacZ Mask, Repo or ElaV Mask, and the intersection were estimated by iterating through each z-slice of the stacks and summing the integrated density values normalized by 255 (assuming 8-bit images for the binary masks).
3. Mask Inversion and Fluorescence Image Processing. All mask images were inverted across the stack to ensure correct masking behaviour during fluorescence processing. Fluorescence signals were isolated within each mask by performing a “Transparent-zero create stack” operation followed by subtraction of the original mask from the resulting stack. This was done for each mask-fluorescence image pair and for the intersection mask with both fluorescence images.
4. Intensity and Mean Fluorescence Quantification (This was not collected for the puc measurement). For each masked fluorescence image, the macro iterated through all z-slices to calculate the sum of normalized integrated density (IntDen/Area) (also from the Image Calculator tool) and mean gray value (Mean) for each slice. These values were accumulated across the stack to compute the total intensity and average mean intensity per ROI.

The following measurements were reported:

– Total and mean fluorescence intensity for Puc-lacZ Mask
– Total and mean fluorescence intensity for Repo or ElaV Mask
– Total and mean fluorescence intensity of Puc-lacZ in regions positive for both masks (Puc-lacZ Mask ∩ ElaV or Repo Mask).

% of Puc-LacZ glia or neurons ware calculated by dividing the total volume of “Puc and Repo or ElaV Mask” by the total volume of Repo or ElaV Mask.

Mean Puc-LacZ intensity in glia or neurons was directly obtained from mean fluorescence intensity of Puc-lacZ in regions positive for both masks (Puc-lacZ Mask ∩ ElaV or Repo Mask).

### Quantification of apoptotic cells

Cells marked by Dcp-1 or GC3Ai were manually counted throughout the z-stack. The distance between apoptotic tumour cells was measured by counting the smallest number of nuclei in-between.

### Statistical tests and experimental reproducibility

Statistical tests used for each experiment are stated in the figure legends. All statistical tests were performed using GraphPad Prism 10.3.1 except for linear correlation (Fig. 3F-G). No samples were excluded except when mentioned in the methods or figure legends (removal of outliers). No sample-size calculations were performed *a priori*. Sample sizes were determined by the number of processed samples (around 5-8 staged adult VNCs per experiment following previous standards (*61–63*)).

For comparing two conditions, an unpaired Student t-test was used when they passed the normality test (D’Agostino and Pearson test) and Mann-Whitney U test when they did not pass the normality test. For comparing more than two conditions, a one-way ANOVA with Tukey’s multiple comparisons test was used on normal data and a Kruskal–Wallis H test with Dunn’s multiple comparisons test for data which did not pass normality.

The correlations between the glia and tumour volumes were tested using a linear model: the glia volume was used as the dependent variable and both condition and tumour volume as explanatory variables, as well as their interaction. n/N numbers are indicated on the figure and their correspondence in the figure legends.

### Data representation

All images displayed are representative. Box plots show minimal value (bottom whisker), first quartile (25th percentile, lower limit of the box), a median of the interquartile range (middle horizontal line), third quartile (75th percentile, the upper limit of the box) and maximal value (top whisker). Individual values are superimposed.

For dot plots, the identity of the dots is described in the legends of the figure and the horizontal line corresponds to the median. Significant and noteworthy p-values are displayed directly on the graph.

## Acknowledgments

We thank members of the Spéder lab for helpful discussion and suggestions; R. Levayer for critical reading of the manuscript; H. Varet of the Biostatistics and Bioinformatics Hub for the analysis of linear correlations; J.C. Luna Escalante for writing the Python code for volume quantification; O. Marshall for help with the analysis of TaDa-PolII datasets. We are very grateful to A. Perez Garijo and H. Steller for the kind gifts of the *QUAS-reaper*, *QUAS-p35* and *QUAS-bsk^DN^ Drosophila* transgenic lines. We thank the VDRC, BDSC and DSHB stock centers for reagents.

## Funding

This work has been funded by recurrent funding from the LabEx Revive (ANR-10-LABX-0073), a Tremplin ERC grant from Agence Nationale de la Recherche (SiStemNiC, ANR-22-ERCC-0005) to PS, an exploratory grant from the Brown Foundation and a Labellisation ARC from the Association pour la Recherche contre le Cancer to PS. This work was supported by a government grant managed by the Agence Nationale de la Recherche under the France 2030 program, with the reference number ANR-24-EXCI-0001, ANR-24-EXCI-0002, ANR-24-EXCI-0003, ANR-24-EXCI-0004, ANR-24-EXCI-0005. MG was supported by a 4-year PhD fellowship from La Ligue Nationale Contre le Cancer. SJ was supported by a 3-year postdoctoral fellowship from Institut Pasteur (Projet Explore) to PS. MH was supported by a 3-year engineer contract from the Association pour la Recherche contre le Cancer to PS.

For the purpose of open access, the author has applied a CC-BY public copyright licence to any Author Manuscript version arising from this submission.

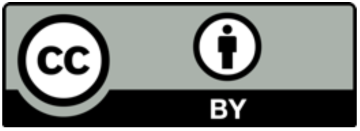

## Author Contributions

Conceptualization: MG, PS

Methodology: MG, CM, PS. CM provided expertise on the tumour model.

Investigation: MG, SJ, MH. MG performed experiments of Figures 1; S1; 2A-C; S2; 3; S3; 4; S4; 5A-D and 6. SJ performed experiments of Figures 1I; 2D-F; S4; 5F-J; S5 and 6. MH helped with experiments of Figures 1I; 2D-F; S4; 5F-J; S5 and 6.

Formal Analysis: DM performed the analysis of the TaDa-PolII datasets. Visualization: MG, SJ, PS

Funding acquisition: PS

Project administration: PS Supervision: PS

Writing – original draft: MG, PS

Writing – review & editing: MG, CM, PS

## Data and materials availability

The TaDa-PolII dataset is available as Datasets S1 and S2. The scripts for volume quantification will be accessible on Zenodo (10.5281/zenodo.16733817). Quantifications are available in the Data source spreadsheet. *Drosophila* transgenic lines made for this study are available upon request to the corresponding author and will be deposited in BDSC. The image datasets generated during and/or analysed during the current study are available from the corresponding author on reasonable request.

## Supplemental information

Supplemental Figures S1–S5 and legends

**Dataset S1.** Excel file containing the differential expression analysis for the TaDa experiments. The columns contain the different outpit avlues from NOISseq analysis.

**Dataset S2.** Excel file containing the Gene enrichment analysis for the TaDa experiments.

Table S1. *Drosophila* genotypes, crosses and culture regimens per experiment Table S2. Tumour growth rates Data Source

