## Supplementary Material for "Cell competition overcomes host tissue resistance to unleash tumour growth in a *Drosophila* brain cancer model"

*Gualtieri et al*

#### Table S1. Genotypes, crosses and regimens

| Figure | Genotypes and crosses | Regimen |
|---|---|---|
| **1B-C, H-I**<br>**S1A** | *UAS- pros$^{RNAi}$, UAS-dicer2/CyO, tub-GAL80; poxn-GAL4/TM6,Tb*<br>*x Nrv2::GFP* | 25°C from embryogenesis until desired time of dissection |
| **1D, F** | *Nrv2::GFP/CyO*<br>*x w1118* | 25°C from embryogenesis until desired time of dissection |
| **1K**<br>**S1B, D** | *Nrv2::GFP, UAS-Raeppli-NLS 53D; act-frt-stop-frt-gal4*<br>*x UAS- pros$^{RNAi}$, UAS-dicer2/CyO, tub-GAL80; poxn-GAL4/TM6,Tb* | 25°C from embryogenesis until desired time of dissection |
| **S1C** | *UAS-Raeppli-NLS 53D; act-frt-stop-frt-gal4*<br>*x UAS- pros$^{RNAi}$, UAS-dicer2/CyO, tub-GAL80; poxn-GAL4/TM6,Tb* | 25°C from embryogenesis until desired time of dissection |
| **2A-C**<br>**S4C-D** | *UAS-prosRNAi, UAS-dicer2/CyO*<br>*x poxn-GAL4/TM6,Tb* | 25°C from embryogenesis until desired time of dissection |
| **2D-F** | *Nrv2::GFP, cyp4g14-QF2; QUAS-His ::mCherry*<br>*x UAS- pros$^{RNAi}$, UAS-dicer2/CyO, tub-GAL80; poxn-GAL4/TM6,Tb*<br>*x poxn-GAL4/TM6,Tb* | 25°C from embryogenesis until desired time of dissection |
| **3A-D** | *actin-LexA, LexAop-GC3Ai*<br>*x UAS- pros$^{RNAi}$, UAS-dicer2/CyO, tub-GAL80; poxn-GAL4/TM6,Tb*<br>*x poxn-GAL4/TM6,Tb* | 25°C from embryogenesis until desired time of dissection |
| **3F-G**<br>**S2** | *UAS- pros$^{RNAi}$, UAS-dicer2/CyO, tub-GAL80; poxn-GAL4/TM6,Tb*<br>*x UAS-syp$^{RNAi}$*<br>*x w1118* | 25°C from embryogenesis until desired time of dissection |
| **4A-C** | *cyp4g15-FRT-STOP-FRT-QF2; poxn-GAL4/TM6,Tb*<br>*x yw, hs-FLP; UAS- pros$^{RNAi}$, UAS-dicer2/CyO, tub-GAL80; QUAS-reaper* | • 25°C from embryogenesis<br>• heatshock 4 h at 37°C at Day 1 OR No heatshock for control<br>• 25°C until desired time of dissection |
| **4D-F** | *cyp4g15-FRT-STOP-FRT-QF2; poxn-GAL4/TM6,Tb*<br>*x yw, hs-FLP; UAS- pros$^{RNAi}$, UAS-dicer2/CyO, tub-GAL80; QUAS-p35* | • 25°C from embryogenesis<br>• heatshock 4 h at 37°C at Day 1 OR No heatshock for control<br>• 25°C until desired time of dissection |
| **5A-E**<br>**S3** | **Tumour**<br>*cyp4g15-FRT-STOP-FRT-QF2; poxn-GAL4/TM6,Tb*<br>*x yw, hs-FLP; UAS- pros$^{RNAi}$, UAS-dicer2/CyO, tub-GAL80; QUAS-Dam*<br>*x yw, hs-FLP; UAS- pros$^{RNAi}$, UAS-dicer2/CyO, tub-GAL80; QUAS-Dam-PolII*<br>**Control**<br>*cyp4g15-FRT-STOP-FRT-QF2; poxn-GAL4/TM6,Tb*<br>*x yw, hs-FLP; ; QUAS-Dam*<br>*x yw, hs-FLP; ; QUAS-Dam-PolII* | • 25°C from embryogenesis<br>• heatshock 4 h at 37°C :<br>- at Day 1 (0 h) OR Day 6 (0 h)<br>• 16 h at 25°C until processing |
| **5F-J** | *puc-LacZ*<br>*x UAS-pros$^{RNAi}$, UAS-dicer2/CyO, tub-GAL80; poxn-GAL4/TM6,Tb*<br>*x poxn-GAL4/TM6,Tb* | 25°C from embryogenesis until desired time of dissection |
| **6**<br>**S4A-B** | *cyp4g15-FRT-STOP-FRT-QF2; poxn-GAL4/TM6,Tb*<br>*x yw, hs-FLP; UAS-pros$^{RNAi}$, UAS-dicer2/CyO, tub-GAL80; QUAS-bsk$^{DN}$* | • 25°C from embryogenesis<br>• heatshock 4 h at 37°C at Day 1 OR No heatshock for control<br>• 25°C until desired time of dissection |
| **S5** | *Nrv2::GFP, tub-GAL80ts; cyp4g15-GAL5*<br>*x UAS-bsk$^{DN}$*<br>*x w1118* | • 18°C until adult eclosion<br>• 29°C until desired time of dissection |

**Table S2. Growth rates of pros$^{poxn}$ tumours without and with JNK inhibition in the cortex glia**

| Genotype | Day | Tumour Volume (μm³) | Calculs |
|---|---|---|---|
| *poxn > pros$^{RNAi}$, cyp > -* | D6 | 351449.031 | |
| | | 299532.688 | |
| | | 415255.406 | |
| | | 514844.375 | |
| | | 358122.906 | |
| | | 341621.5 | |
| | | 550357.062 | |
| | | 290634.69 | Mean |
| | | 539969.81 | 406865.2742 |
| | D9 | 781915.938 | |
| | | 842715.719 | |
| | | 1043912.812 | |
| | | 716402.406 | |
| | | 847978 | |
| | | 612403.56 | |
| | | 1180989 | |
| | | 1102869.88 | Mean |
| | | 679455.375 | 867626.9656 |
| | | **Growth Ratio** | **2.13** |
| | | **% Growth** | **113.25** |

| Genotype | Day | Tumour Volume (μm³) | Calculs |
|---|---|---|---|
| *poxn > pros$^{RNAi}$, cyp > bsk$^{DN}$* | D6 | 388599 | |
| | | 558731.5 | |
| | | 458716.656 | |
| | | 444376.938 | |
| | | 775272.375 | |
| | | 567134.75 | |
| | | 466352.781 | |
| | | 701863.75 | Mean |
| | | 451517.844 | 534729.5104 |
| | D9 | 899631.438 | |
| | | 418725.09 | |
| | | 583401.125 | |
| | | 507532.19 | |
| | | 555471.312 | |
| | | 862030.562 | |
| | | 673336.062 | Mean |
| | | 556318.812 | 632055.8239 |
| | | **Growth Ratio** | **1.18** |
| | | **% Growth** | **18.20** |

### Supplemental Figures and Captions for

**Cell competition overcomes host tissue resistance to unleash tumour growth in a *Drosophila* brain cancer model**

*Gualtieri et al*

#### Figure S1

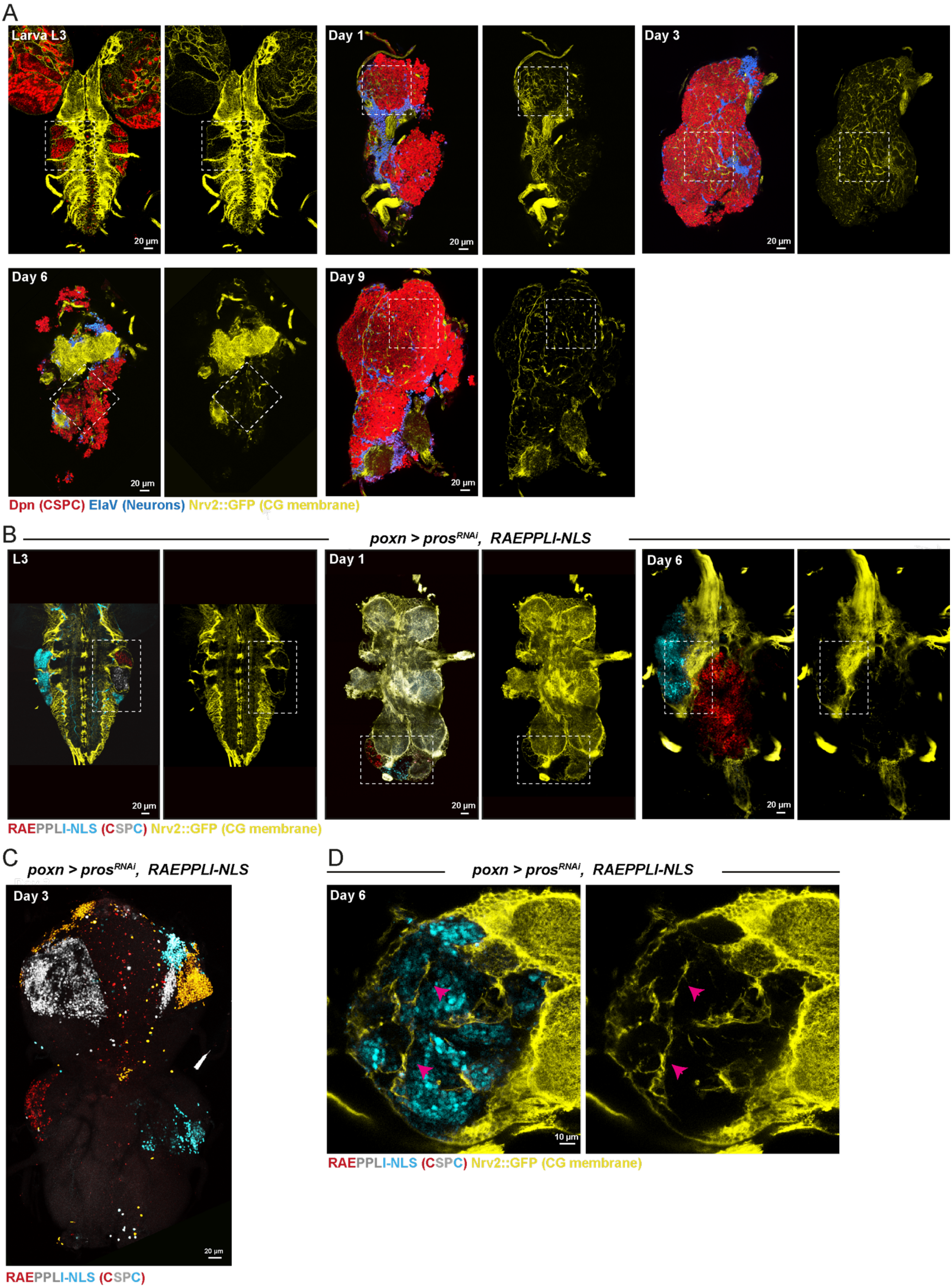
A
Larva L3
Day 1
Day 3
Day 6
Day 9
20 µm
Dpn (CSPC) ElaV (Neurons) Nrv2::GFP (CG membrane)
B
poxn > pros^RNAi, RAEPPLI-NLS
L3
Day 1
Day 6
20 µm
RAEPPLI-NLS (CSPC) Nrv2::GFP (CG membrane)
C
poxn > pros^RNAi, RAEPPLI-NLS
Day 3
20 µm
RAEPPLI-NLS (CSPC)
D
poxn > pros^RNAi, RAEPPLI-NLS
Day 6
10 µm
RAEPPLI-NLS (CSPC) Nrv2::GFP (CG membrane)

**Figure S1. Cancer stem cell-driven tumours remodel the host glial microenvironment, related to Figure 1**

(**A**) Confocal images of whole VNCs illustrating the progressive remodelling of CG membranes upon tumour progression, from larval to adult stage. Dashed white squares indicate the cropped areas shown in the close-up images of Fig. 1H.

(**B**) Confocal images of whole VNCs where *pros*$^{poxn}$ tumours have been labelled by Raeppli-NLS induced at larval stage, at third larval stage (L3) and days 1 and 6 of adulthood. CG membrane is marked by Nrv2::GFP (yellow). Dashed white rectangles indicate the cropped areas corresponding to the close-up images in Fig. 1K.

(**C**) Projection of a confocal image showing a whole VNC at day 3 in which *pros*$^{poxn}$ tumours have been labelled by Raeppli-NLS induced at larval stage.

See also Table S1.

14 **Figure S2**

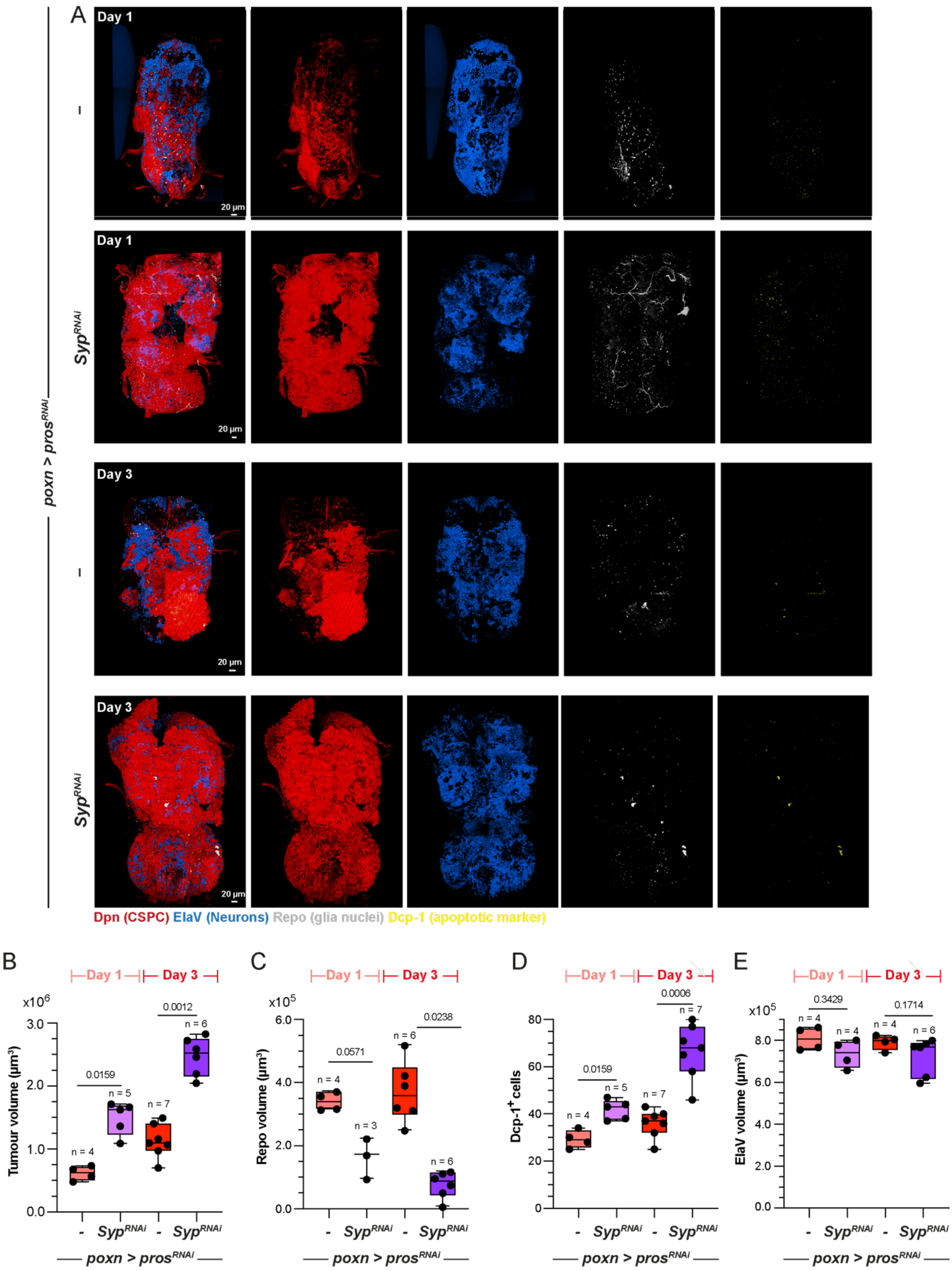

A
Day 1
Day 3
–
Syp^RNAi
poxn > pros^RNAi
20 µm
Dpn (CSPC) ElaV (Neurons) Repo (glia nuclei) Dcp-1 (apoptotic marker)
B
C
D
E
Day 1
Day 3
Tumour volume (µm³)
x10^6
Repo volume (µm³)
x10^5
Dcp-1+ cells
ElaV volume (µm³)
0.0159
0.0012
0.0571
0.0238
0.0006
0.3429
0.1714
n = 4
n = 5
n = 7
n = 6
n = 3

**Figure S2. Fast-growing tumours enriched in cancer stem cells cause increased glial loss, related to Figure 3**

(**A**) 3D reconstruction of confocal images showing whole VNCs staining for tumour (Dpn, red), glial cells (Repo, gray), neurons (ElaV, blue) and apoptotic cells (Dcp-1, yellow) for control $pros^{poxn}$ tumours ($poxn$ > $pros^{RNAi}$) and tumours in in which Syncrip (*syp*), a factor essential for the generation of more restricted progenitors with lower proliferative capacity, has been knocked down, resulting in larger tumours ($poxn$ > $pros^{RNAi}$, $syp^{RNAi}$) at days 1 and 3 of adulthood.

(**B**) Box plot of tumour (Dpn) volume for $pros^{poxn}$ and $Syp^{RNAi}$ $pros^{poxn}$ tumours at days 1 and 3 of adulthood. n = VNC. Mann-Whitney U tests for each day.

(**C**) Box plot of glia (Repo) volume per VNC for $pros^{poxn}$ and $Syp^{RNAi}$ $pros^{poxn}$ tumours at days 1 and 3 of adulthood. n = VNC. Mann-Whitney U tests for each day.

(**D**) Box plot of apoptotic cell (Dcp-1) numbers per VNC for $pros^{poxn}$ and $Syp^{RNAi}$ $pros^{poxn}$ tumours at days 1 and 3 of adulthood. n = VNC. Mann-Whitney U tests for each day.

(**E**) Box plot of neuron (ElaV) volume per VNC for $pros^{poxn}$ and $Syp^{RNAi}$ $pros^{poxn}$ tumours at days 1 and 3 of adulthood. n = VNC. Mann-Whitney U tests for each day.

For box plots, individual values are superimposed.

See also Table S1.

**Figure S3**

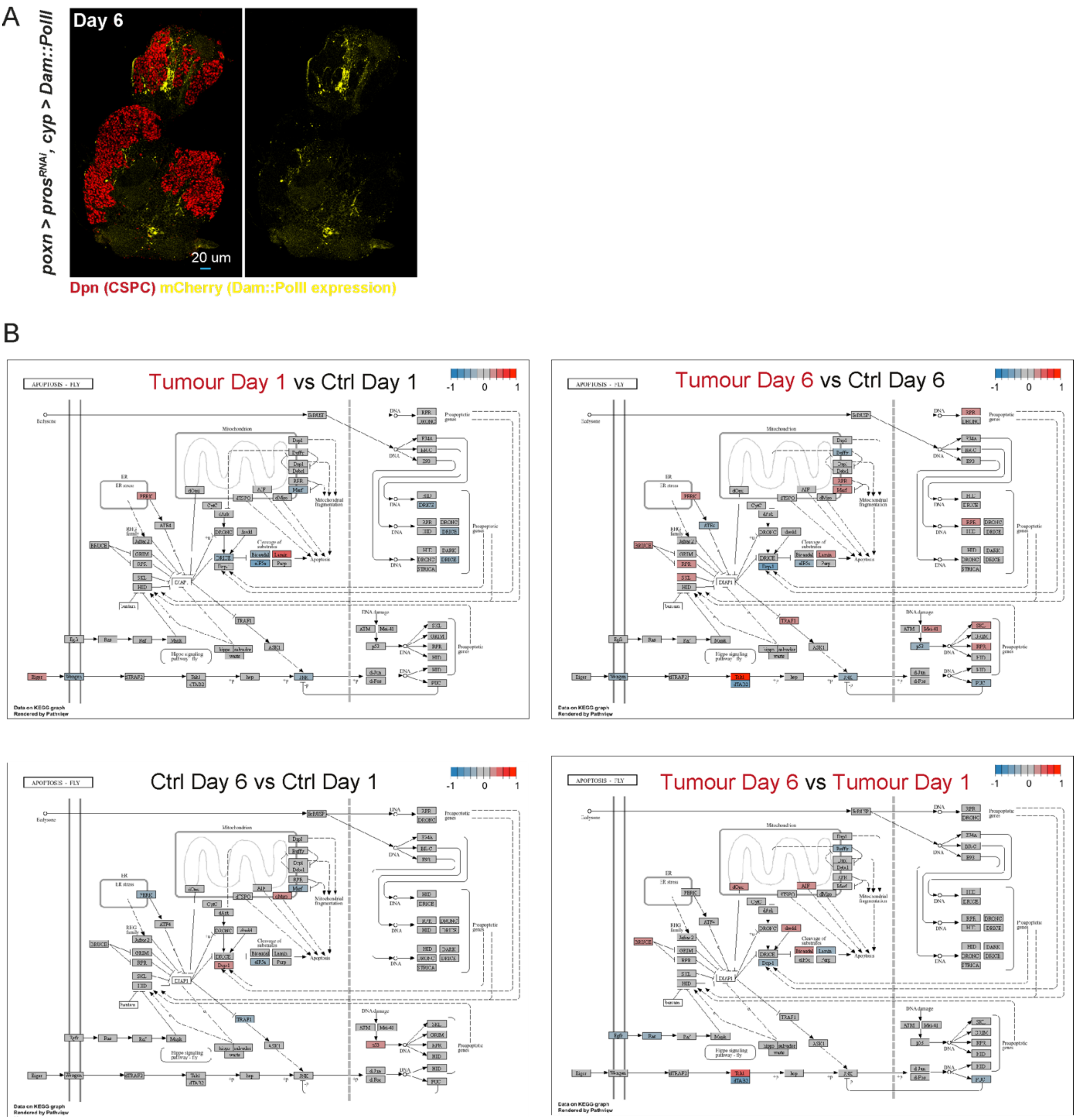
A
poxn > prosRNAi, cyp > Dam::PolII
Day 6
20 um
Dpn (CSPC) mCherry (Dam::PolII expression)
B
APOPTOSIS - FLY
Tumour Day 1 vs Ctrl Day 1
Tumour Day 6 vs Ctrl Day 6
Ctrl Day 6 vs Ctrl Day 1
Tumour Day 6 vs Tumour Day 1
-1 0 1
Data on KEGG graph
Rendered by Pathview

**Figure S3. Transcriptional analysis of cortex glia along tumour progression reveals a downregulation of the JNK pathway, related to Figure 5**

(**A**) Confocal image showing the expression of Dam-PolII (*QUAS-Dam-PolII* construct) specifically in the cortex glia (*cyp4g15-QF2* driver line) under the control of the Q system in a tumour context (*poxn* > $pros^{RNAi}$, *cyp* > *Dam-PolII*) at day 6.

(**B**) Differential expression heatmap for genes of the KEGG Drosophila apoptosis (dme04214) pathway for four comparisons: $pros^{poxn}$ tumour Day 1 versus Control (no tumour) Day 1; $pros^{poxn}$ tumour Day 6 versus Control Day 6; Control Day 1 versus Control Day 6; $pros^{poxn}$ tumour Day 1 versus $pros^{poxn}$ tumour Day 6. Fold change scale is Log2.

See also Table S1 and Datasets 1 and 2.

#### Figure S4

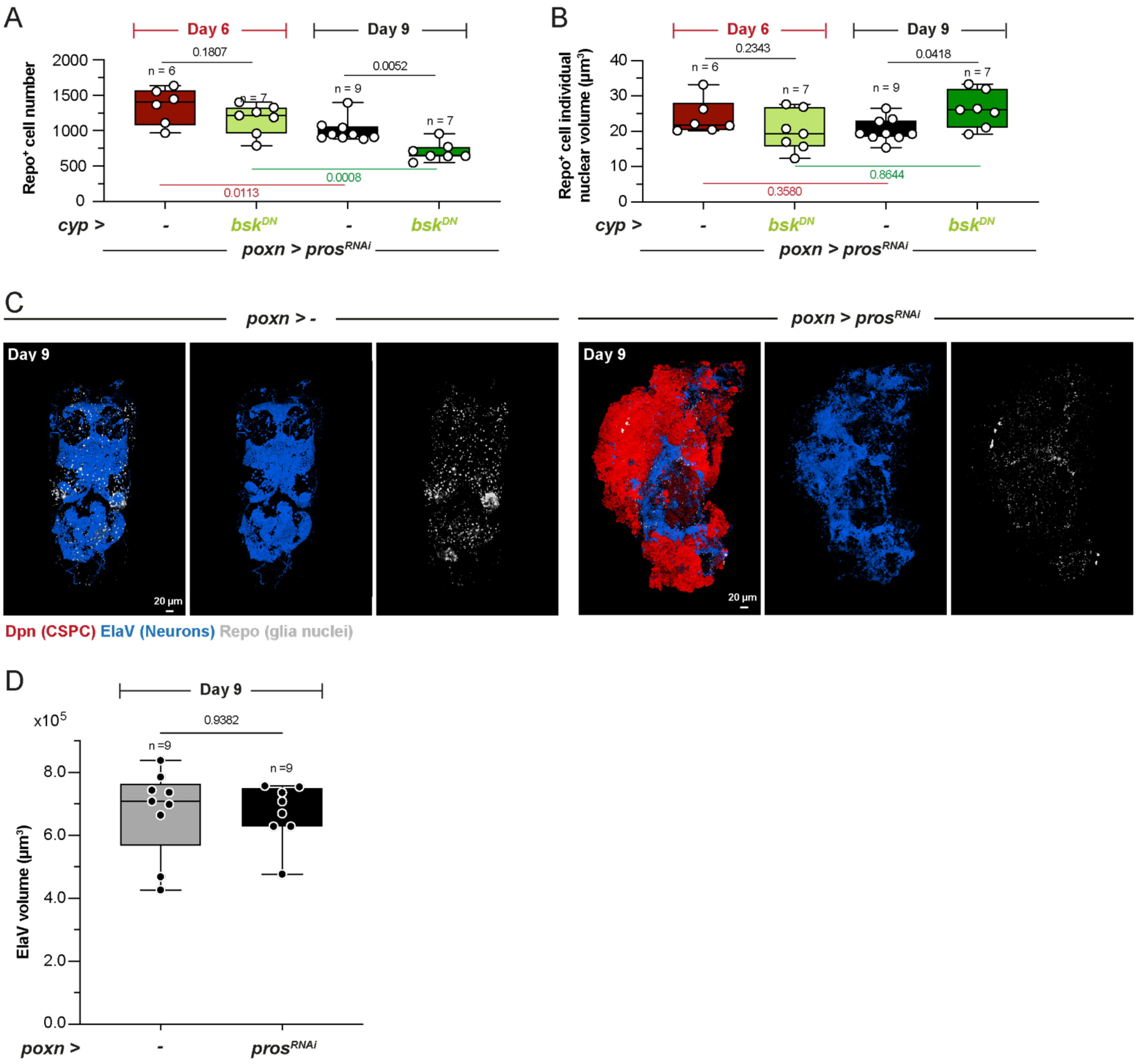
A
Day 6
Day 9
Repo+ cell number
n = 6
n = 7
n = 9
n = 7
0.1807
0.0052
0.0008
0.0113
cyp >
-
bskDN
-
bskDN
poxn > prosRNAi
B
Day 6
Day 9
Repo+ cell individual nuclear volume (µm3)
0.2343
0.0418
0.8644
0.3580
C
poxn > -
poxn > prosRNAi
Day 9
Day 9
20 µm
20 µm
Dpn (CSPC) ElaV (Neurons) Repo (glia nuclei)
D
Day 9
x10^5
0.9382
n =9
n =9
ElaV volume (µm3)
poxn >
-
prosRNAi

**Figure S4. Glial and neuronal loss change in tumours where the JNK pathway is downregulated in the cortex glia, related to Figure 6**

(**A**) Box plot of glia (Repo$^{+}$) numbers per VNC for (*poxn* > *pros*$^{RNAi}$*, cyp* > -) and (*poxn* > *pros*$^{RNAi}$*, cyp* > *bsk*$^{DN}$) at days 6 and 9 of adulthood. n = VNC. Kruskal Wallis with with Dunn's multiple comparisons test between days. Mann-Whitney U tests between control and tumour conditions for each day.

(**B**) Box plot of glia (Repo$^{+}$) individual nuclear volume per VNC for (*poxn* > *pros*$^{RNAi}$*, cyp* > -) and (*poxn* > *pros*$^{RNAi}$*, cyp* > *bsk*$^{DN}$) at days 6 and 9 of adulthood. n = VNC. Kruskal Wallis with with Dunn's multiple comparisons test between days. Mann-Whitney U tests between control and tumour conditions for each day.

(**C**) 3D reconstruction of confocal images showing whole VNCs stained for neuronal (ElaV, blue) and glial (Repo, gray) nuclei for control (*poxn* > -) and *pros*$^{poxn}$ tumour (*poxn* > *pros*$^{RNAi}$) conditions at day 9 of adulthood.

(**D**) Box plot of total neuron (ElaV$^{+}$) volume for control (*poxn* > -) and *pros*$^{poxn}$ tumour (*poxn* > *pros*$^{RNAi}$) conditions at day 9 of adulthood. n = VNC. Student's t-test.

See also Table S1.

**Figure S5**

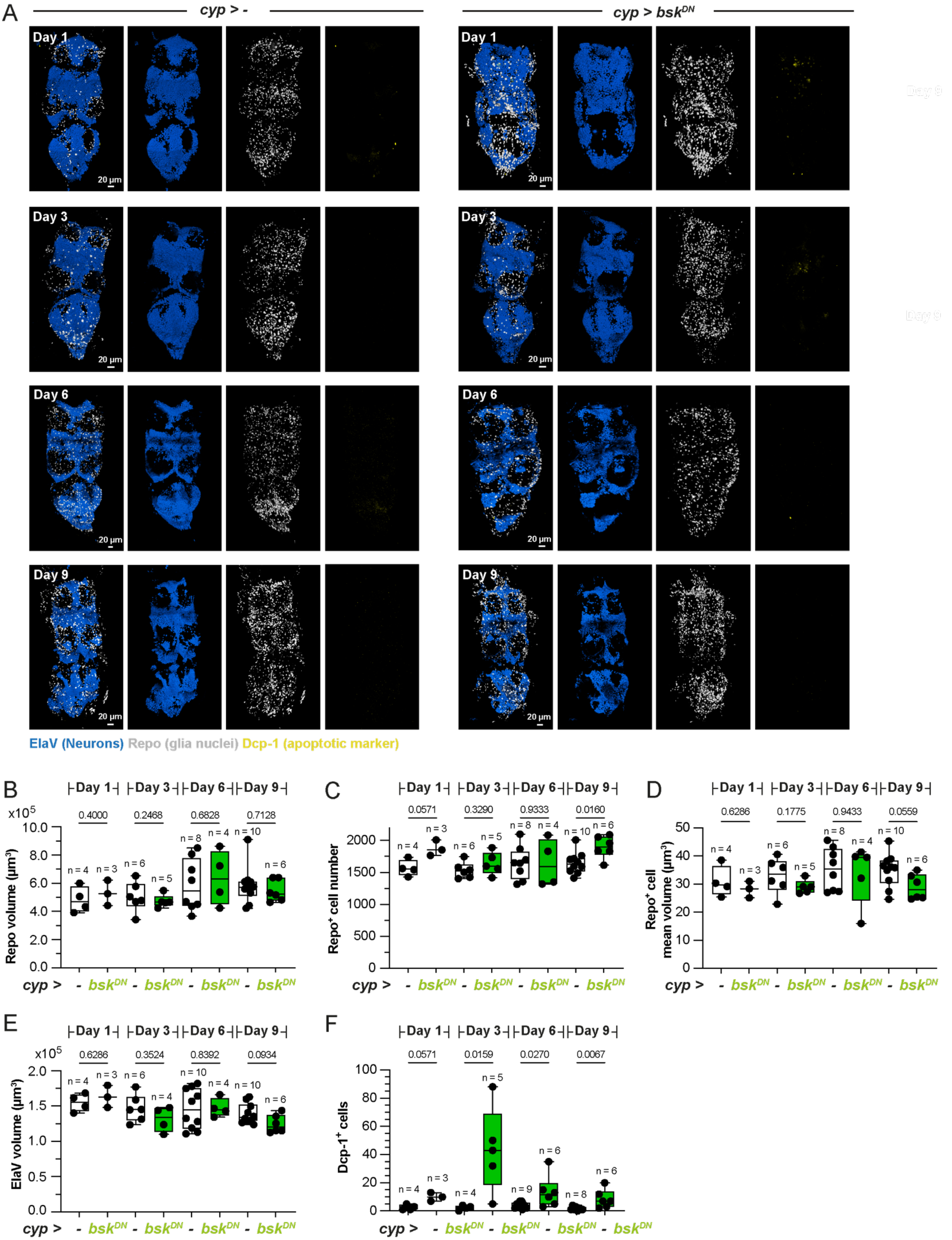
A
cyp > -
cyp > bsk^DN
Day 1
Day 3
Day 6
Day 9
20 µm
ElaV (Neurons) Repo (glia nuclei) Dcp-1 (apoptotic marker)
B
Day 1 Day 3 Day 6 Day 9
0.4000 0.2468 0.6828 0.7128
Repo volume (µm³)
x10^5
cyp > - bsk^DN - bsk^DN - bsk^DN - bsk^DN
C
0.0571 0.3290 0.9333 0.0160
Repo+ cell number
D
0.6286 0.1775 0.9433 0.0559
Repo+ cell mean volume (µm³)
E
0.6286 0.3524 0.8392 0.0934
ElaV volume (µm³)
F
0.0571 0.0159 0.0270 0.0067
Dcp-1+ cells

**Figure S5. The JNK pathway plays a protective role towards neurons in normal conditions, related to Figure 6**

(**A**) 3D reconstruction of confocal images showing whole VNCs staining for glial cells (Repo, gray), neurons (ElaV, blue) and apoptotic cells (Dcp-1, yellow) in control condition (*cyp* > -) and during inhibition of the JNK pathway in CG (*cyp* > $bsk^{DN}$) for days 1, 3, 6 and 9 of adulthood. To prevent $bsk^{DN}$ expression during embryonic and larval stages, the TARGET system was used (see Methods), resulting in a switch from 18°C to 29°C at day 0 of adulthood. As such, days 1, 3, 6 and 9 are at 29°C (and not 25°C as used previously), which correspond to around two days later than at 25°C.

(**B**) Box plot of glia (Repo) number for (*cyp* > -) and (*cyp* > $bsk^{DN}$) at days 1, 3, 6 and 9 of adulthood (at 29°C). n = VNC. Mann-Whitney U tests for each day.

(**C**) Box plot of glia (Repo) volume per VNC for (*cyp* > -) and (*cyp* > $bsk^{DN}$) at days 1, 3, 6 and 9 of adulthood (at 29°C). n = VNC. Mann-Whitney U tests for each day.

(**D**) Box plot of neuron (ElaV) volume per VNC for (*cyp* > -) and (*cyp* > $bsk^{DN}$) at days 1, 3, 6 and 9 of adulthood (at 29°C). n = VNC. Mann-Whitney U tests for each day.

(**E**) Box plot of apoptotic cell (Dcp-1) numbers per VNC for (*cyp* > -) and (*cyp* > $bsk^{DN}$) at days 1, 3, 6 and 9 of adulthood (at 29°C). n = VNC. Mann-Whitney U tests for each day.

For box plots, individual values are superimposed.
See also Table S1.
